# Metabolic-imaging of human glioblastoma live tumors: a new precision-medicine approach to predict tumor treatment response early

**DOI:** 10.1101/2022.04.12.487679

**Authors:** Mariangela Morelli, Francesca Lessi, Serena Barachini, Romano Liotti, Nicola Montemurro, Paolo Perrini, Orazio Santo Santonocito, Carlo Gambacciani, Matija Snuderl, Francesco Pieri, Filippo Aquila, Azzurra Farnesi, Antonio Giuseppe Naccarato, Paolo Viacava, Francesco Cardarelli, Gianmarco Ferri, Paul Mulholland, Diego Ottaviani, Fabiola Paiar, Gaetano Liberti, Francesco Pasqualetti, Michele Menicagli, Paolo Aretini, Giovanni Signore, Sara Franceschi, Chiara Maria Mazzanti

**Author notes:** Competing interests: The authors have no relevant financial or non-financial interests to disclose.

## Abstract

**Background:** Glioblastoma (GB) is the most severe form of brain cancer, with a 12-15 month median survival. Surgical resection, temozolomide (TMZ) treatment, and radiotherapy (RT) remain the primary therapeutic options for GB, and no new therapies have been introduced in recent years. This therapeutic standstill is primarily due to preclinical approaches that do not fully respect the complexity of GB cell biology and fail to test efficiently anti-cancer treatments. Therefore, better treatment screening approaches are needed. In this study, we have developed a novel functional precision medicine approach to test the response to anticancer treatments in organoids derived from the resected tumors of glioblastoma patients.

**Methods:** GB organoids were grown for a short period of time to prevent any genetic and morphological evolution and divergence from the tumor of origin. We chose metabolic imaging by NAD(P)H fluorescence lifetime imaging microscopy (FLIM) to predict early and non-invasively ex-vivo anti-cancer treatment responses of GB organoids. TMZ was used as the benchmark drug to validate the approach. Whole-transcriptome and whole-exome analyses were then performed to characterize tumor cases stratification.

**Results:** Our functional precision medicine approach was completed within one week after surgery and two groups of TMZ Responder and Non Responder tumors were identified. FLIM-based metabolic tumor stratification was well-reflected at the molecular level, confirming the validity of our approach, highlighting also new target genes associated with TMZ treatment and identifying a new 17 gene molecular signature associated with survival. The number of promoter methylated tumors for the MGMT gene was higher in the responsive group, as expected, however, some non-methylated tumor cases turned out to be nevertheless responsive to TMZ, suggesting that our procedure could be synergistic with the classical MGMT methylation biomarker.

**Conclusions:** For the first time, FLIM-based metabolic imaging was used on ex-vivo live glioblastoma organoids. Unlike other approaches, ex-vivo patient-tailored drug response is performed at an early stage of tumor culturing with no animal involvement and with minimal tampering with the original tumor cytoarchitecture. This functional precision medicine approach can be exploited in a range of clinical and laboratory settings to improve the clinical management of GB patients and implemented on other cancers as well.

## Introduction

Glioblastoma (GB) is the most common malignant primary brain tumor. Overall, the prognosis of patients with this disease is poor, with a 12-15 month median survival^1^. Because of the diffuse aggressive nature of GB cell invasion into the brain parenchyma, no GB patient has been cured to date^2^. As illustrated by the vast number of drugs and therapeutic strategies under investigation for the treatment of GB, there is a major effort to develop more effective therapies to treat this highly malignant and therapy-insensitive disease. Unfortunately, the success of these new therapies has been rather disappointing: maximal resection, and temozolomide (TMZ) treatment and radiotherapy (RT) remain the best option for quite several years^3^. GB standard-of-care TMZ is a DNA-alkylating agent discovered in the 1970s and approved by the FDA in 2005^3^. Responsive patients have the O6-methylguanine DNA methyltransferase (MGMT) gene with a methylated promoter and show higher survival rates than patients with a hypomethylated MGMT gene. Despite its low specificity, the MGMT promoter status represents the only clinical biomarker available to predict for TMZ response^4^.

In this glioblastoma context, effective treatment options and biomarkers of drug response are a major unmet medical requirement. For several years, conventional monolayer cell cultures have been widely used to test drug efficiency, although the lack of tissue architecture and complex characteristics of these models fails to recapitulate the true biological processes in vivo. Recent advances in organoid technology have revolutionized in vitro culture tools for biomedical research by creating powerful 3D-dimensional models, that better preserve the local cytoarchitecture and native cell-cell interactions of original tumors^5^. Despite numerous ex-vivo drug testing approaches leverage on 3d-in vitro GB models, they still have shortcomings and none fully captures the complexity of each individual glioblastoma cellular organization and composition overlooking therefore the relevance of how the tumor microenvironment affects tumor behavior and drug response^6 7 8^. These drug testing approaches have several limiting requirements such as: a) extended time of performance that leads to a long period of in vitro tumor culturing with a consequent molecular and morphological transformation diverging from the parental tumor in vivo^6^; b) technical measurements requiring dissociation of the original tumoral tissue down to a single cell suspension, losing therefore the tumor cytoarchitecture and cell-cell interaction characteristics^8^; c) use of non-human animal models which implies laborsome procedures and introduction of biases due to host organism-tumor interactions^9^.

To address these limitations it is necessary to have a treatment-testing approach that doesn’t need an extended in vitro tumor culturing, that is non-invasive to minimize tumor tampering, that uses a biomarker of response that is precocious and anticipates early enough tumor behavior so to be applied at an early stage of in vitro tumor culturing and no animal involvement. Furthermore, because of the highly aggressive progression of the disease, an overall rapid test and selection of the optimal drug regimen for individual glioblastoma patients is crucial and a method to predict the drug response, early, without wasting patients’ lifetime, before the onset of the therapy could be transformative for GB patients.

Here in this study, to answer these requirements, we offer a unique novel treatment-testing approach in patient-derived glioblastoma 3D organoids. We first developed a protocol to generate an in vitro vital patient-derived IDH1/2 wild-type GB 3D model that we termed “glioblastoma explant” (GB-EXP), which, unlike other models ^6 7 8^, is minimally handled, briefly grown in culture with no animal involvement, no dissociation and passaging to preserve the parental cytoarchitecture. To build a treatment-response predictor tool, we applied an imaging method called FLIM that exploits the intrinsic auto-fluorescence molecular properties of NAD(P)H, a metabolic enzymatic cofactor, that is associated with the metabolic state of the cell/tissue. Cancer cell metabolic status is known to be an early predictor of cellular behavior in response to a treatment ^10 11 12^. The intracellular metabolic cofactor NAD(P)H (reduced form of nicotinamide adenine dinucleotide) may be protein-bound or protein-free in the cell, and these states affect its fluorescence decay, with bound NAD(P)H typically exhibiting longer lifetimes than free NAD(P)H. In cancer, metabolism shifts were investigated, and authors reported an increase in NAD(P)H fluorescence lifetimes (increase of NAD(P)H bound/free ratio) as cells become less proliferative ^13^, showing drug responsiveness after treatment. FLIM measures NAD(P)H lifetimes, representing a powerful non-invasive tool to monitor, in real time, metabolic activities in living cells and tissues ^10 12^. NAD(P)H-FLIM has the advantage of being a fast and non-invasive method that can be applied to in vitro cancer organoids at an early stage of in vitro culturing without interfering with tumor viability and structure, avoiding divergence from parental tumor. To support this approach as an effective method for evaluating the response to anti-cancer treatments, we used TMZ as our benchmark drug since it is the only approved drug used in GB, and extensive knowledge has been achieved at the molecular level. We achieved the classification of the cases into TMZ responsive and non-responsive tumors. This stratification performed out solely using our NAD(P)H -FLIM approach on live GB organoids in vitro, was then correctly corroborated by conventional drug testing assays and by next generation sequencing analyses. The TMZ responsive and non-responsive groups were statistically significantly distinguished at the genomic and transcriptomic level confirming the accuracy of our approach.

This novel approach in the assessment of GB treatment outcome, could be exploited as a tool for improving patient-tailored therapeutic strategies by testing single of combinations of drugs, and new treatments. It can also be used for large-scale screening of new pharmaceutical compounds and implemented on other tumors as well.

## Materials and Methods

### Human glioblastoma tissue collection

The study was performed in accordance with the Declaration of Helsinki and the sample collection protocol was approved by the Ethics Committee of the University Hospital of Pisa (787/2015). Tumors were obtained from patients who had undergone surgical resection of histologically confirmed GB after obtaining informed consent. Samples were obtained from the Neurosurgery Department of the “Azienda Ospedaliero-Universitaria Pisana” or from the Unit of Neurosurgery of Livorno Civil Hospital. Sixteen male and female patients were included in the present study. All patients were diagnosed with GB with no previous history of brain neoplasia and did not carry R132 IDH1 or R172 IDH2 mutations. In five of the 16 patients, neurosurgeons were able to collect the core and periphery of the tumor with the help of neuronavigation–guided microsurgical techniques. Tumor samples at the periphery were first obtained when GB was identified during surgery, whereas tumor samples at the core were obtained from the resected tumor mass. When the tumor had a large area of central necrosis, the tumor located outside the necrotic area was selected. The patients clinical and demographic data are presented in Table 1S. Surgically resected tumors were collected and stored in MACS tissue storage solution (Miltenyi Biotec, Bergisch Gladbach, Germany) at 4?C for 2–4 h. All patient-derived surgical GB tissues were de-identified before processing. See supplementary materials for more information.

### Glioblastoma cell line spheroid cultures

Spheroids were generated from T98G and U87 GBM cell lines using the hanging drop method, as previously described ^14^. Once spheroids were formed, they were used to prepare cultures in Matrigel and in suspension. Twenty drops per well were transferred to a 4-well chamber coverglass (Nalge Nunc International) and covered with 300 µL of Matrigel Growth Factor Reduced Basement Membrane Matrix, phenol red-free Matrigel (Corning). After gel solidification for 30’ at 37°C, cell medium without no red phenol, containing 89% of DMEM low glucose (for T98G) and high glucose (for U87), 10% FBS, and 1% penicillin-streptomycin was added. Cultures were placed in a 37°C, 5% CO2, and 90% humidity sterile incubator. The medium was replaced every 72 hours. See the supplementary materials for more information on 2D cell lines and 3D spheroids.

### Glioblastoma organoids/explants cultures

The procedure used to produce explant cultures is shown in Figure. 1a. Fresh GB tumors or frozen samples, after a quick defrosting in a water bath at 37°C, were washed with DPBS in a sterile dish and cut with a scalpel into pieces <1 mm^2^. Samples were then run on a gentleMACS Dissociator (Miltenyi Biotec) to mechanically dissociate them into 80–200 μm macrosuspensions, which were filtered through a 70-micron cell strainer to exclude smaller tissue pieces. Tissue suspensions were placed in coverglass chamber slides (Nalge Nunc International) and then covered with 300 µL of Matrigel or in suspension. Cultures were placed in a 37°C, 5% CO2, and 90% humidity sterile incubator and grown for a maximum of 10/14 days without no passaging. The GB-EXPs were cultured without additional growth factors to reduce tampering. Drug treatment was initiated 3 days after the GB-EXPs were cultured.

**Figure 1.**
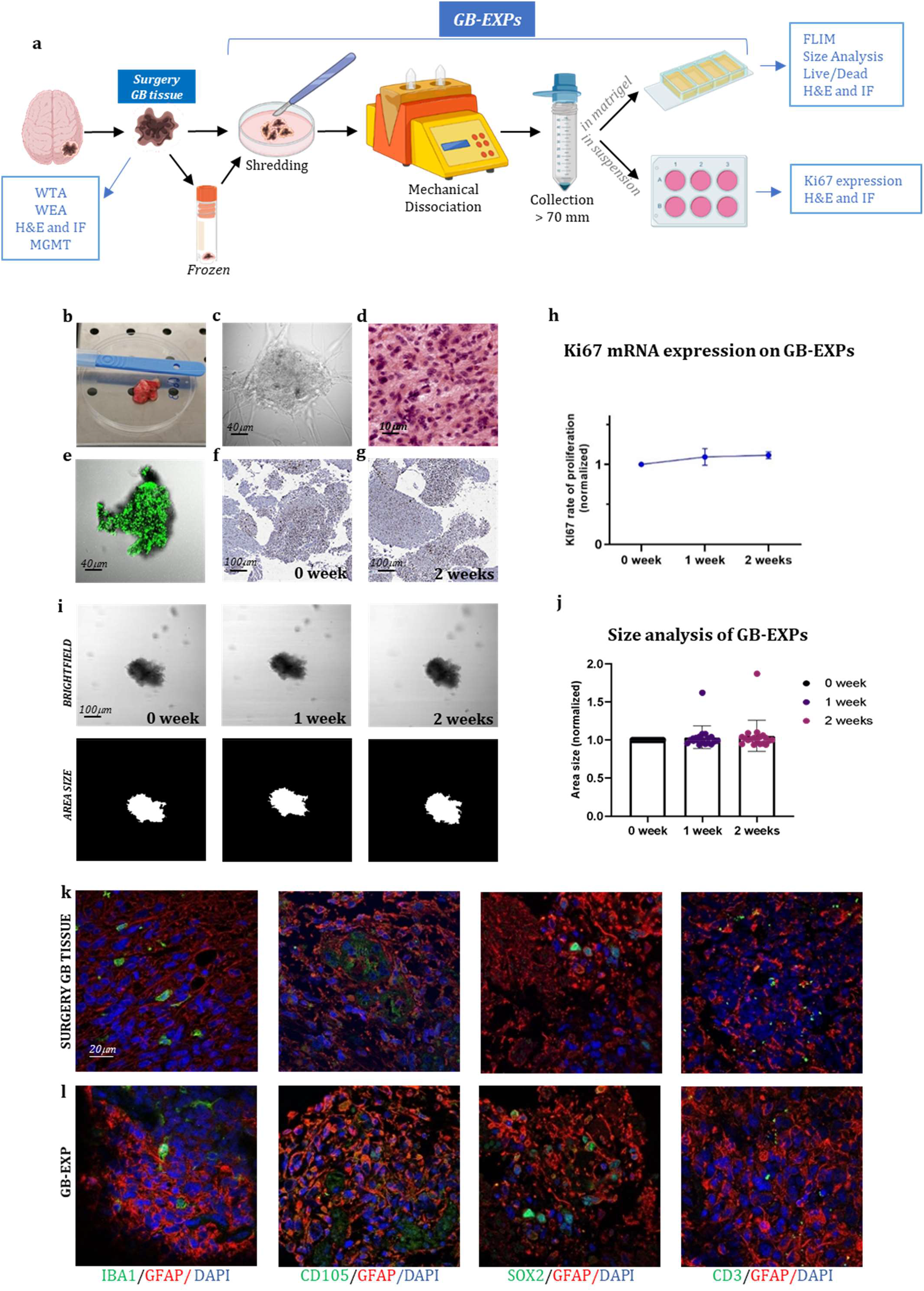
GB-EXPs cultures. (a) Experimental design. (b) A surgery tumor sample. (c) GB-EXP embedded in matrigel. (d) H&E staining of a surgery GB tissue. (e) Live/dead staining of a GB-EXP at 2 weeks after culturing. (f-g) Ki67 immunostaining of GB-EXPs in suspension at 0 week and 2 weeks. (h) Ki67 mRNA expression analysis of 13 GB case-derived GB-EXPs at 0 and 2 weeks. Graph represents mean ± s.d. of triplicated measures. (i) Representative brightfield and area size images of a GB-EXP in matrigel at 0 week, 1 week and 2 weeks. (j) Size analysis 20 GB case-derived GB-EXPs at 0, 1 and 2 weeks. (k-l) Immunofluorescence assays of surgery GB tissue (k) and GB-EXPs (l). GFAP (astrocytes), IBA1 (microglia), SOX2 (stem cells), CD105 (endothelial cells) and CD3 (lymphocytes). **Abbreviations**: WTA, Whole Transcriptome Analysis; WEA, Whole Exome Analysis; FC, Flow Cytometry; H&E, Hematoxylin, and Eosin; MGMT, MGMT promoter methylation analysis.

### TMZ Drug treatments

TMZ (Sigma, St. Louis, MI, USA) was used in this study. TMZ was dissolved in DMSO to prepare a stock concentration of 100 mM and then diluted to the required concentrations with a complete cell culture medium. 2D GB U87 and T98G cell lines were treated at 30% confluence, replacing the medium with fresh medium containing TMZ 100 μM for treated cells or an equal volume of DMSO for controls. Cultures were exposed to TMZ for 24, 48, and 72 h in all experiments. Spheroids from U87 and T98G cell lines, both in Matrigel and in suspension, were treated the day after they were cultured, replacing the medium with fresh medium containing 600 μM of TMZ. For FLIM experiments, spheroids in Matrigel were exposed to TMZ for 24, 48, and 72 h, and for sizing and Ki67 expression analysis for 1 and 2 weeks. GB-EXPs, both in Matrigel and in suspension, 3 days after being cultured, were treated with TMZ three days after culture. The medium was replaced with fresh medium containing 600 μM TMZ for the treated explants or an equal volume of DMSO as a control. GB-EXPs in Matrigel were exposed for 24 h, 48 h, and 72 h for FLIM experiments, whereas GB-EXPs in suspension were treated at 1 week and 2 weeks for sizing, live/dead, and immunofluorescence experiments. Fresh conditioned medium for both the GB-EXPs and spheroid cultures was replaced every 72 h.

### Lifetime Imaging

Fluorescence lifetime imaging was performed with an Olympus Fluoview 3000 confocal microscope using a 405 nm LDH-P-C-375B (Picoquant) excitation laser for NAD(P)H ^15,16^. For the control and treated samples, 6–12 FLIM measurements were acquired. The phasor approach was used to analyze NAD(P)H-FLIM data and was performed using the SimFCS suite as previously described ^17^. Following the instructions reported in *“Two-component analysis of fractional NAD(P)H distribution”* ^17^, we extrapolated from phasor plots the NAD(P)H free/bound fractional distribution curves of the controls and treated samples, creating a mean distribution curve for controls and one for treated samples (Supplementary Figure 1). To find differences between controls and treated NAD(P)H fractional distribution curves, for each of the 125 parts SimFCS dividing the curves, a p-value was calculated using the parametric Student’s t-test, and fractions with p values <0.05 were considered significant. Each of the 125 parts consists of a number of image pixels with a specific NAD(P)H free/bound fraction. Therefore, the curve segmentation implemented by the system reflects different portions/pixels of the tumor image, consequently reporting intra-sample metabolic heterogeneity. We calculated the percentage of response (%DR), considering only the treated distribution, as follows: %DR = ∑SIGNIFICANT AREA/TOTAL AREA. where:1) the SIGNIFICANT AREA is the area obtained by adding the areas of the histograms, resulting in a p-value<0.05, when compared with controls; and 2) the TOTAL AREA is the area obtained by adding the areas of the 125 histograms. The histogram area was calculated by multiplying the base (corresponding to the unit at each x-axis point) by the height (corresponding to the mean of the normalized pixels at each y-axis point). The %DR was calculated after 24, 48, and 72 h of treatment. The final percentage of DR was obtained by calculating the weighted average. Weighted average calculation: The weight given to the average was driven by the time point, given a value of 3, 2, and 1 for 72, 48, and 24 h, respectively. 72hr time had the highest weight since it is the standard time at which cells reach metabolic adaptation ^18^. Stratification of samples was performed using %DR as follows: Non Responder (Non-Resp/NR): %DR<5% Low Responder (LR):5≤%DR<20, Medium Responder (MR):20≤%DR<50, High Responder (HR): %DR>50.

### GB-EXPs Size Analysis

The growth of explants was studied using brightfield images acquired at 0 day, after 1 and 2 weeks after TMZ treatment, using an Olympus Fluoview 3000 microscope and at 20X magnification. A minimum of 14 to a maximum of 81 untreated and control GB-EXPs were imaged at each time point. Different wells were used for GB-EXPs sizing and FLIM. GB-EXPs growth over time was measured using OrganoSeg Software, kindly donated by the Department of Biomedical Engineering, University of Virginia, USA ^19^.

### Histology and stainings

The tissues and explants were fixed for 24 h in 10% neutral-buffered formalin (Sigma-Aldrich) at room temperature. For immunofluorescence, CD105 polyclonal (Thermo Fisher, PA5-94980), CD3 monoclonal (F7.2.38) (Thermo Fisher, MA5-12577), Sox2 polyclonal (Thermo Fisher, 48-1400), GFAP monoclonal (ASTRO6) (Thermo Fisher, MA5-12023), GFAP polyclonal (Abcam, ab7260), Iba1 polyclonal (Wako 091-19741) primary antibodies were then applied at dilutions of 1:400, 1:20, 1:100, 1:100, 1:250, 1:1000, respectively, overnight at 4°C, and visualized using Olympus Fluoview 3000 confocal microscope at a magnification of 60X. See supplementary materials for more information.

### Next-generation sequencing analyses

The RNA-seq library was prepared using Illumina Stranded Total RNA Prep with a Ribo-Zero Plus kit (Illumina). Whole exome library preparation was performed using Illumina DNA Prep with Enrichment (Illumina, San Diego, CA, USA) following the manufacturer’s instructions, starting with 500 ng of DNA. Sequencing was performed on a NextSeq 500 (Illumina, San Diego, CA, USA) with a reading length of 101 bp (Supplementary Materials).

### Statistical analysis

All summary data are presented as means ± s.d.. All statistical analyses were performed using R and GraphPad Prism software (GraphPad 7.0). The sample size (n) values used for statistical analyses are provided in the text and supplementary materials. Individual data points are graphed or can be found in the source data. Tests for differences between two groups were performed using the Student’s two-tailed unpaired t-test, as specified in the figure legends. No data points were excluded from the statistical analyses. Statistical significance was set at p < 0.05. Linear discriminant analysis was performed using JMP10 software (SAS Institute).

## Results

### Tumor Samples

The dataset included 21 patient-derived surgery GB tissues, five of which consisted of the core and periphery of the tumor from the same GB patient, obtained with the help of neuronavigation–guided microsurgical techniques. The patients included 10 men (62%) and six

(38%) women in the age group of 30–80 years. All tumor samples were derived from primary IDH1/2 wild-type glioblastoma samples. Each resected sample was labeled with information on cerebral localization and molecularly characterized for IDH1/2 mutation and MGMT methylation status, as shown in Supplementary Table 1. Furthermore, MGMT methylation analysis revealed MGMT methylation discordance between the core (c) and peripheral (p) portions of samples GB3, GB4, GB6, and GB7, highlighting intra-tumor heterogeneity (Supplementary Table 1). Supplementary Table 2 reports the pathological diagnosis and information about the patients’ therapeutic administrations. The samples were subjected to several analyses, as shown in Supplementary Table 3.

### In vitro culturing of patient-derived glioblastoma explants (GB-EXPs)

We created a vital human GB-patient-derived 3D tumor culture in vitro (GB-EXPs) (Figure. 1a). Twenty-one tumor pieces were first washed with PBS and a specific lysis buffer to remove debris and red blood cells, respectively (Figure 1b). Mechanical dissociation and filtration produced a suspension of tumor pieces ranging in size from 70µm to 200µm (Fig. 1c). Tumor fragments were are rapidly processed to maximize their viability and ensure good explant quality. GB-EXPs were cultured immediately after resection, but also after short time storage at -140?, confirming that viable cultures can be grown either from fresh or flash-frozen DMSO supplemented media, thus facilitating the whole procedure^20^. GB-EXPs were cultured by embedding them into Matrigel or suspension (no single-cell dissociation was performed) (Figure 1c). To reduce clonal selection and maintain tumor heterogeneity and parental cytoarchitecture, explant cultures were not grown for more than 2 weeks. In vitro these GB-EXPs represent the actual pathological conditions in vivo as closely as possible. Before culturing, GB tissues were subjected to H&E staining and histological analysis, which was performed by an expert pathologist to confirm the GB features (Figure 1d, Supplementary Figure 2). GB-EXPs still showed active proliferation within 2 weeks of culture (Supplementary Figure 3). Moreover, the vitality of GB-EXPs cultured in Matrigel was determined by setting up overnight live-imaging analyses, which revealed an intensive cell activity particularly evident at the surface of the explant in contact with the surrounding cells and neighboring explants, as shown in Supplementary Video 1. To evaluate the viability of GB-EXPs, we performed a live/dead cell viability assay (Figure 1e) and Ki67 immunohistochemistry (Figure 1f,g). In GB-EXPs, Ki67 expression analysis by immunohistochemistry showed positivity at either 0 or 2 weeks of culture, as shown in Figure 1f,g. Furthermore, the rate of proliferation was explored using Ki67 mRNA expression analysis (Figure 1h) and size analysis ^19^ (Figure1i,j), which showed a slight gradual increase in the proliferation rate over time, specifically from week 0 to week 2, confirming the results of other culturing approaches reported in the literature ^6^. To assess whether GB-EXPs maintain the cyto-composition of parental tumors, we further characterized and explored cellular diversity among surgical GB tissues and GB-EXPs by choosing a panel of GB markers including GFAP (astrocytes), IBA1 (microglia), SOX2 (stem cells), CD105 (endothelial cells), and CD3 (lymphocytes). The presence of these cell types was confirmed by immunofluorescence assays on GB-EXPs, indicating retention of vasculature features and lymphocytes after two weeks of in vitro culturing (Figure 1k,l).

### FLIM metabolic-imaging approach validation in known glioblastoma *in vitro* systems

Initially, to validate the efficacy of our FLIM-based metabolic imaging approach, we evaluated it using GB U87 and T98G cell lines, known to be TMZ-responsive and non-responsive, respectively, ^21^. The two cell lines were tested in the 2D and 3D systems.

#### 2D glioblastoma cell lines system

We investigated two different glioblastoma commercial cell lines that are sensitive and resistant to TMZ treatment, U87, and T98G ^21–23^. To monitor intracellular molecular changes associated with drug treatment, FLIM image data were recorded from 12 fields of view for each slide of both cell lines 72hr post treatment, targeting the autofluorescence of the intracellular metabolic cofactor NAD(P)H.

Figure 2a-c shows representative images of a cellular field of 2D-U87 and 2D-T98G cells, including brightfield images (top row) and phasor-FLIM NAD(P)H lifetime map (on the bottom) colored in accordance with the color bar defined on the side. The color bar defines the metabolic pathway from NAD(P)H in the bound state (red/magenta) to NAD(P)H in the free state (green/white), as explained in the Materials and Methods section (Supplementary Fig. 1a-b). Based on phasor-based FLIM data analysis, we obtained a fractional NAD(P)H distribution curve of free and bound NAD(P)H molecules for each image (see Materials and Methods and Supplementary Fig. 1a-b), which identifies a metabolic signature that goes from an oxidative phosphorylation phenotype with low free/bound NAD(P)H fractions to a glycolytic phenotype with high free/bound NAD(P)H ^17^. In Figure 2b and d, we report the fractional NAD(P)H mean distribution curves of the control and TMZ-treated 2D-U87 (Fig. 2b) and 2D-T98G cell lines (Fig. 2d). The average distribution curves were broken down into different histograms, representing each specific fraction of free-state NAD(P)H and consequently of protein-bound molecules (see Materials and Methods). Differences between fractional NAD(P)H mean distribution curves were evaluated by statistically comparing each histogram of the treated group against the same one in the control group using Student’s t-test (Fig. 2b,d) (see Materials and Methods). Therefore, the difference between the two curves was expressed as a percentage of drug response (%DR) calculated considering the area of the significant histograms (in green, Figure 2b,d) out of the total histogram area under the treated GB-EXPs curve (see Materials and Methods).

**Figure 2:**
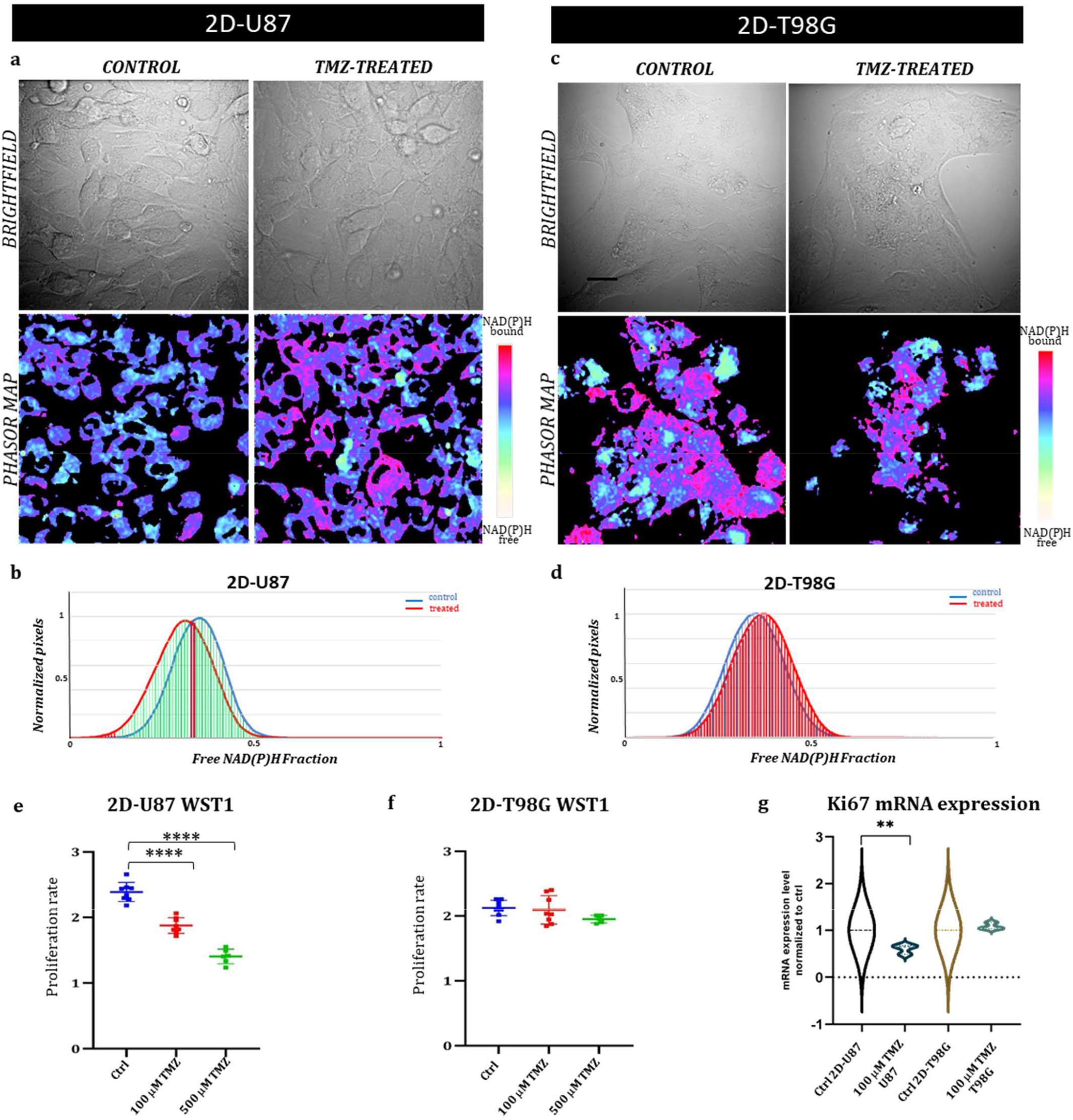
2D glioblastoma cell lines system: FLIM-based metabolic imaging in GB commercial cell lines TMZ responder 2D-U87 and TMZ non responder 2D-T98G. (a,c) Representative images of 2D-U87 (a) and 2D-T98G (c) subdivided in brightfield images and the corresponding phasor maps. Scale bar, 30 ?m. (b,d) NAD(P)H fractional mean distribution curves of control (blue) and treated cells (red) for 2D-U87 (b) and 2D-T98G cells (d), 72 hrs post TMZ treatment. (e,f) Proliferation curves of 2D-U87 (e) and 2D-T98G cell lines (f) after 72 hrs (p<0.001, Student’s t test). (g) Ki67 mRNA expression using real-time PCR in control and treated cells, 72hrs after treatment (p=0,002, Student’s t-test).

In TMZ-treated sensitive 2D-U87 cells, distinctive FLIM signatures were observed with a statistically significant left-bound shift (red curve) towards a higher fraction of bound NAD(P)H compared with TMZ-treated cells to control cells, with 92%DR (Fig. 2b). The 2D-T98G cell lines showed no changes in FLIM signatures, as represented by the overlapping of the control and treated distributions (0%DR) (Fig. 2d).

The percentages of differences between the NAD(P)H fractional distribution curves of the control and treated samples for both cell lines were well reflected, as well as in the phasor maps obtained for each cellular field, as shown in Fig. 2a,c.

We linked these shifts in FLIM data distribution curves after treatment to changes in NAD(P)H lifetimes, as reported in several studies in the literature^12^. The observed shift towards higher fractions of bound NAD(P)H is often associated with a more oxidative-oriented metabolism, which is characteristic of less proliferative cells^11,12,24,25^. This is consistent with the results of the proliferation assay shown in fig 2e. Figures 2e and 2f show the proliferation assay for both cell lines 72 h post-treatment with 100µM and 500µM µM TMZ. The drug doses were chosen based on previous reports^21–23^. 2D-U87 cells showed a statistically significant decrease in proliferation with both TMZ dosages (Fig.2e), while 2D-T98G cells showed no statistical difference between treated and control cells (Fig. 2f).

As an indicator of cell proliferation, we also measured by real time PCR the expression level of Ki67 at 72 h in TMZ-treated and control 2D-U87 and 2D-T98G cell lines (Fig.2g). 2D-U87 TMZ-treated cell lines showed a statistically significant reduction in Ki67 mRNA expression compared to control cells, consistent with the observed lower rate of cell proliferation, while no difference was detected in 2D-T98G cells (Fig. 2g).

#### 3D glioblastoma spheroids system

To mimic the 3D structures of GB-EXPs, TMZ responder 3D-U87 and TMZ Non Responder 3D-T98G spheroids were used to measure the impact of TMZ drug response on FLIM metabolic imaging data. In the literature, several TMZ dilutions ranging from 1µM to 1mM have been studied on GB organoids ^26^ with the most effective doses between 250µM and 1mM. In line with these results, we selected 600µM. FLIM data were acquired at 72 h for a total of 10 spheroids with characteristic dimensions in the 70-200 µm range for each experimental condition. In Figure 3a,c, representative images of a spheroid are shown, including brightfield images (top row) and NAD(P)H-FLIM phasor map (bottom) colored in accordance with the color bar defined on the side. As reported for the 2D cell line system, in Figure 3b,d, we obtained the fractional NAD(P)H mean distribution curves of the control and TMZ-treated 3D-U87 (Figure 3b) and 3D-T98G spheroids (Figure 3d). After 72 h of TMZ exposure in responsive 3D-U87 spheroids, the treated fractional NAD(P)H mean curve showed a statistically significant shift towards higher fractions of NAD(P) H-bound molecules when compared to controls (see Figure 3b), as demonstrated by a 55%DR. As previously discussed, this indicates an increase in the oxidative metabolism typical of a less proliferative state ^27 10 12^. The %DR is shown by green histograms in Figure 3b and is reflected in the phasor map of a representative spheroid, as shown in Figure 3a,c.

**Figure 3.**
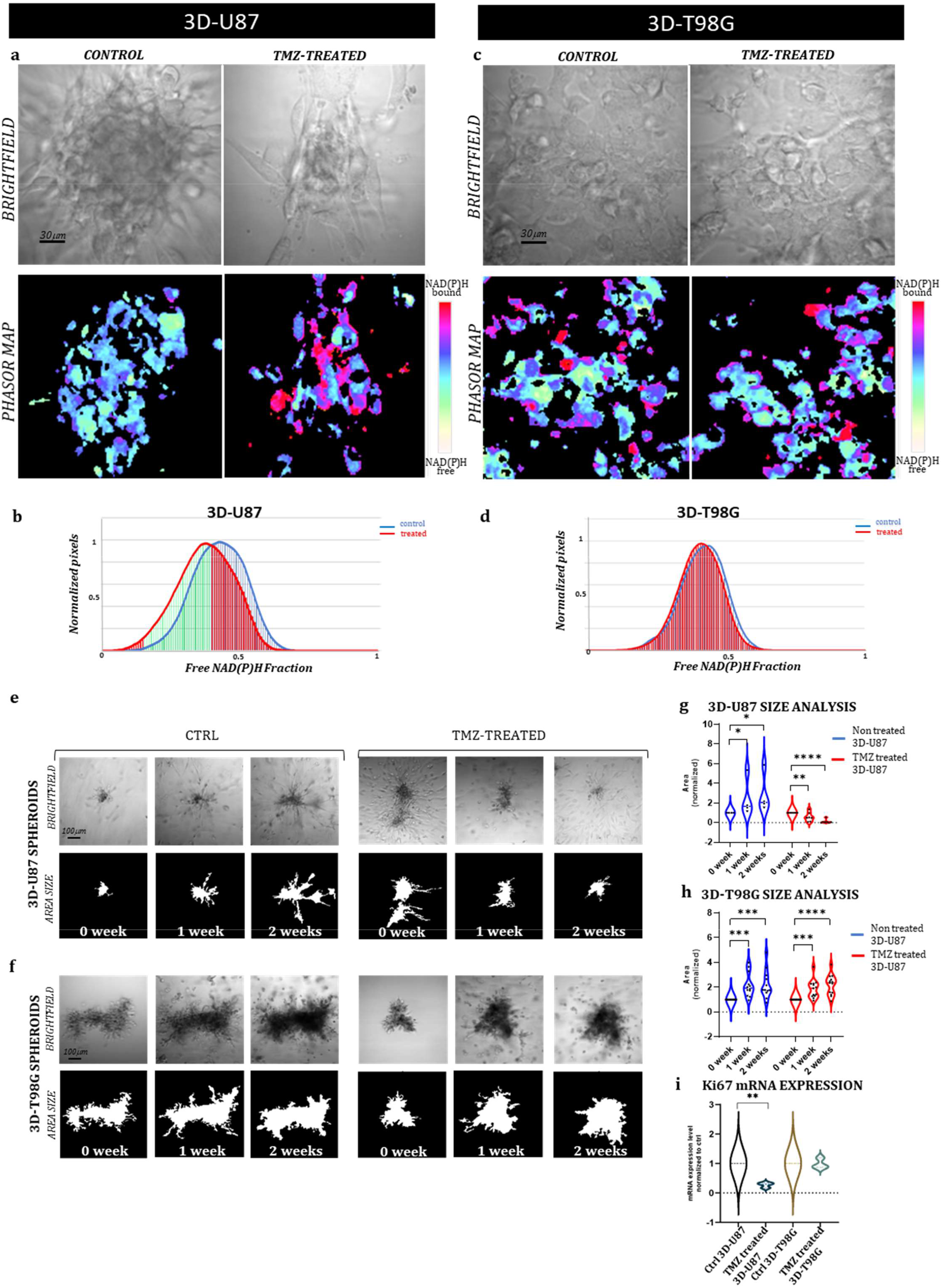
FLIM-based metabolic imaging in 3D-U87 and 3D-T98G. (a,c) Representative images of 3D-U87 (a) and 3D-T98G (c) controls and treated cells, subdivided in brightfield images of a spheroid and the corresponding phasor map displayed in a false color scale with the coding shown by the bar on the side of the figure. Scale bar, 30 μm. (b,d) NAD(P)H fractional mean distribution curves of controls (blue) and treated cells (red) of 3D-U87 (b) and 3D-T98G cells (d). (b) 3D-U87 cells show a statistically significant higher fraction of bound-state NAD(P)H in the treated group compared to the ctrl group (55%DR). (d) In 3D-T98G no difference was found between ctrl and treated cells in fractions of NAD(P)H bound molecules (0%DR). (e,f) Representative brightfield, and area size images of a 3D-U87 (e) and a 3D-T98G (f) control and TMZ 600 μM treated sample at 0 week, 1 week and 2 weeks. Scale bar, 100 μm. (g) Size analysis of 3D-U87 cells reveals at 1 and 2 weeks a significant increased area for controls (n=12) (pvalue= 0.03, Student’s t test), and a significant decreased area for TMZ treated samples (n=12) (p-value=0.003 and p-value<0.00002, respectively, Student’s t test). (h) Size analysis of 3D-T98G cells reveals at 1 and 2 weeks a significant increased area for controls (n=12) (p-value=0.0001, Student’s t-test), and a significant increased area for TMZ treated samples (n=12) (pvalue= 0.001, p-value<0.00002, respectively, Student’s t test). 1 and 2 week areas are normalized to spheroids area at 0 week. (i) Ki67 mRNA expression at 72hrs shows a statistical decrease in rate of proliferation in 3D-U87 TMZ treated compared to controls (p-value=0.002. Student’s t test), unlike 3D-T98G.

To support the FLIM results, we performed spheroid size measurements over time using dedicated software (OrganoSeg) ^19^. Control and TMZ-treated spheroids for both cell lines were followed up for two weeks after treatment and photographed at 0, 1, and 2 weeks. A representative bright-field image of 3D-U87 and 3D-T98G spheroids at each time point is shown in Figure 3e,f together with the matched area size image for both experimental conditions (control and treated). The area size measurement of the 3D-U87 images at each time point revealed a statistically significant increase in size for the control spheroids (n=12) (p-value=0.03, Student’s t-test), and as expected, a statistically significant decrease was detected for the TMZ-treated spheroids (n=12) (1week, p-value=0.003 and 2 weeks p-value<0.00002, respectively, Student’s t-test) (Figure 3g). In 3D-T98G cells, a statistically significant increase in size was recorded at 1 and 2 weeks for both control (n=12) (p =0.0001, Student’s t-test) and TMZ-treated spheroids (n=12) (p =0.001, p <0.00002, respectively, Student’s t-test), confirming their resistance to TMZ treatment (Figure 3h). Ki67 real-time expression analysis at 2 weeks after TMZ treatment confirmed the FLIM readouts and size results (p =0.002. Student’s t-test) (Fig 3i)..

### FLIM metabolic imaging of TMZ treated patient-derived GB-EXPs

Using the methodological criteria set by the supporting technical experiments on GB cell lines, we assessed the TMZ response in 21 glioblastoma patient-derived tumors. For each patient, GB-EXPs with similar dimensions between 70 and 200µm for drug testing, and a minimum of 14 to a maximum of 33 explants were analyzed (Table 4S). The GB-EXPs of the tumor case in Figure 4a show overlapping fractional NAD(P)H mean distribution curves between the treated (red) and control (blue) explants, which is indicative of an identical distribution of bound and free NAD(P)H^12^ (Figure 4b) with a 0%DR. In Figure 4c and d, we report examples of GB-EXPs derived from the GB15 case, resulting in distinctive phasor maps. NAD(P)H fractional mean distribution curves were also distinguishable between the control and treated cases. The distribution curves (Figure 4e,f) showed a statistically significant shift of the red curves towards more abundant bound NAD(P)H molecular species in TMZ-treated explants compared to the control ones, with 59.3%DR (Figure 4e) and 90.9%DR (Figure 4f), corresponding to 24 and 72 h TMZ treatment, respectively (Table 5S). Larger amounts of bound-state NAD(P)H reflect oxidative metabolism, which is typical of less proliferative cells ^13 10^ and therefore a responsive tumor.

**Figure 4.**
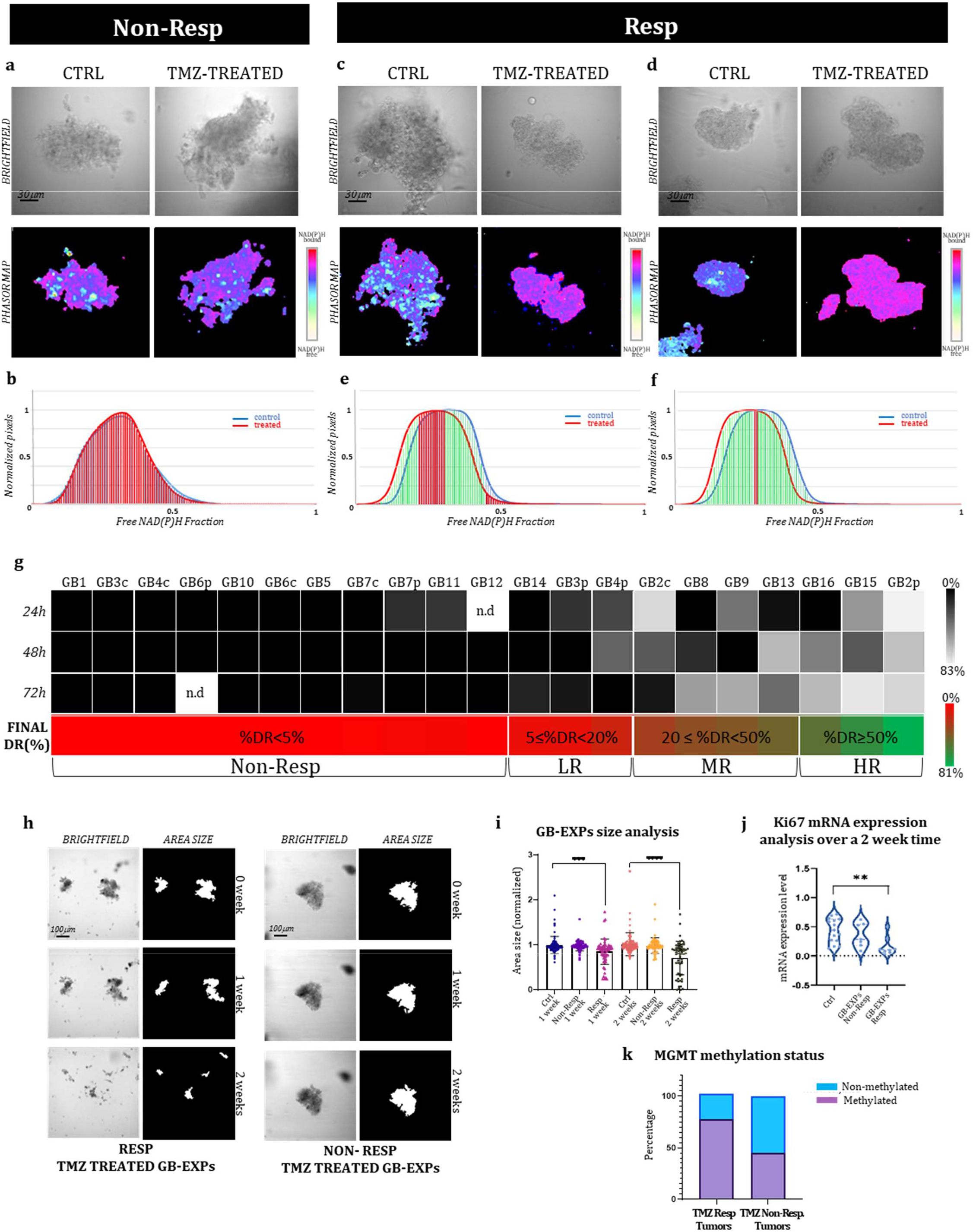
FLIM-based metabolic imaging on GB-EXPs. (a,c,d) 72 hr post-treatment one Non-Resp and two Resp tumor derived GB-EXPs are shown including a brightfield image and the corresponding phasor map. (b) In the Non-Resp case (a) NAD(P)H fractional mean distribution curves (72hrs) overlap between control (blue) and treated GB-EXPs (red) (b). (e,f) NAD(P)H fractional mean distribution curves show a left-bound shift of the red curves (treated GB-EXPs). (g) %DR at 24, 48, and 72 hr, represented by a grey scale is shown for each GB case. The annotation of Resp and Non-Resp was assigned on the basis of the Final %DR, obtained from the weighted average of the 3 time points (shown with a green-red color bar for each GB case). Samples are further subdivided based on Final %DR into several categories. (h) Representative brightfield and area size images of a Resp and of a Non-Resp case-derived GB-EXP in matrigel at 0, 1 and 2 weeks. (i) Size analysis of Controls, Resp and Non-Resp patients-derived GB-EXPs (n=215, n=130, n=124 respectively) at 1 and 2 weeks. A statistically significant decrease at 1 and 2 weeks (p=0.0002; p<0.0001, respectively; Student’s t test) is shown for the Resp group. (j) Ki67 mRNA expression using ddPCR in Resp (n=8) and Non Resp (n=7) cases after 2 weeks of TMZ treatment (*p=0.003, Student’s t test). (k) MGMT promoter methylation status in Resp (n=9) and Non Resp (n=11) cases.

Overall, to assess the final annotation of TMZ Responder (Resp) or Non Responder (Non-Resp), we calculated for each case a Final %DR to TMZ treatment (see Materials and Methods), as shown in Figure 4g (green-red color scale), leading us to stratify our tumor samples into 11 Non-Resp (%DR<5) and 10 Resp. The Resp group was further subdivided into several categories: Low Responders (LR) (5≤%DR<20), Medium Responders (MR) (20≤%DR<50), and High Responders (HR) (%DR≥50) (Figure 4g). It is noteworthy to point out that between core and peripheral portions of the 5 GB tumor cases included in the dataset, we observed a different drug response behavior (Supplementary Figure 4). As expected, GB2, GB3, and GB4 core regions showed a more drug-resistant behavior compared to peripheral tumor portions ^28^ in particular, GB2c was assessed as MR and GB2p as HR, while GB3c/4c were NR and GB3p/4p were LR. GB6c/p and GB7c/p were both assessed as Non-Resp (Figure 4g).

To corroborate the FLIM-based metabolic imaging drug efficacy predictions in stratifying samples in Resp and Non-Resp, we performed a size analysis on all 21 sample-derived GB-EXPs. Images were acquired at 0, 1, and 2 weeks for both control and TMZ-treated cases for a minimum of 15 to a maximum of 81 GB-EXPs per sample (Table 4S). The examples reported in Figure 3h show a clear reduction in size for the Resp GB-EXP compared to the Non-Resp. Overall, the area measurement of the GB-EXPs Resp group revealed a statistically significant decrease at 1 and 2 weeks after TMZ treatment (p=0.0002 and p<0.0001, respectively; Student’s t-test), thus supporting the FLIM-based metabolic imaging predictions (Figure 4i).

To further support our results, we measured the changes in Ki67 mRNA expression 2 weeks after TMZ treatment. Differential analysis of Ki67 expression was performed on seven Non-Resp and eight Resp samples. Ki67 mRNA expression was significantly reduced in the Resp group after TMZ treatment compared to that in the controls, indicating a lower proliferation rate (p=0.003, Student’s t-test). (Figure 4j). No significant difference was found in the Non-Resp group.

In accordance with what has been reported in the clinic and in the literature ^29^, we observed a higher percentage of methylated cases (77%) in the Resp tumor group than in the Non-resp tumor group (45%) (Figure 4k).

### Genome-wide analyses

We performed next-generation sequencing analyses to characterize the genetic background of the Resp and Non-Resp tumors to establish a correlation between the TMZ response phenotype identified by our FLIM approach and the underlying molecular profile.

#### Whole Transcriptome Analysis (WTA)

WTA was performed on nine Resp and nine Non-Resp samples, including the cases provided with a core and a peripheral portion. Differential expression analysis identified 42 statistically significant genes between the two experimental groups (Table 6S). In Figure 5a, a heatmap analysis was initially run excluding the peripheral tumor portions to avoid repeated measures derived from similar genetic backgrounds with their core counterparts. Results showed that seven Resp and seven Non-Resp samples perfectly clustered on the basis of the 42 gene expression levels, supporting that distinct TMZ responsive and non-responsive phenotypes are well reflected at the molecular level. Among the 42 genes (Table 6S), we identified several genes involved in the TMZ response that are worth mentioning. The EGFR gene, which showed a significant downregulation in Non-Resp samples, is consistent with literature data showing that glioblastoma TMZ-resistant cell lines lack EGFR activation and expression^30^. Our results are, as well, confirmed for the upregulation of the CA9 gene in the Non-Resp group, the inhibition of which enhances the sensitivity of glioma cells to TMZ treatment, and highlights the value of developing small molecules or antibodies against the CA9 pathway, for combination therapy with TMZ ^31^(Table 9S, Fig 5a). In the Non-Resp group, we identified downregulation of the FGFR3 gene, which is associated with a poor response to TMZ treatment^32^. The same authors reported that a combination treatment of vinblastine (VBL) and mebendazole (MBZ) with TMZ was more effective in reducing the cell number when glioblastoma cells had low expression levels of FGFR3. Most noteworthy are the genes that we report in Fig. 5b, BIRC3 and NDRG2, which show differential gene expression that varies gradually according to %DR, supporting phasor-FLIM drug response stratification. Some studies in 2016 and 2021^33 34^ report that BIRC3 gene was found expressed at a higher level in recurrent GB than in newly diagnosed GB and emerged as a novel driver of TMZ therapeutic resistance, suggesting that, during TMZ therapy, concurrent BIRC3-specific inhibition could be exploited for enhanced benefit. NDRG2 showed an opposite trend compared to BIRC3 (Fig. 5b). The main mechanism underlying NDRG2 silencing in gliomas remains unknown. There is also debate on whether NDRG2 gene activity reflects the survival of glioma patient ^35^. Furthermore, we identified two genes that were completely silenced in all samples of the Non-Resp group, ANKRD28 and PTPRD (Figure 5a,b). While the ANKRD28 gene has an unknown role in GB, the tyrosine phosphatase PTPRD is a tumor suppressor that is frequently inactivated and mutated in GB and other human cancers ^36^.

**Figure 5.**
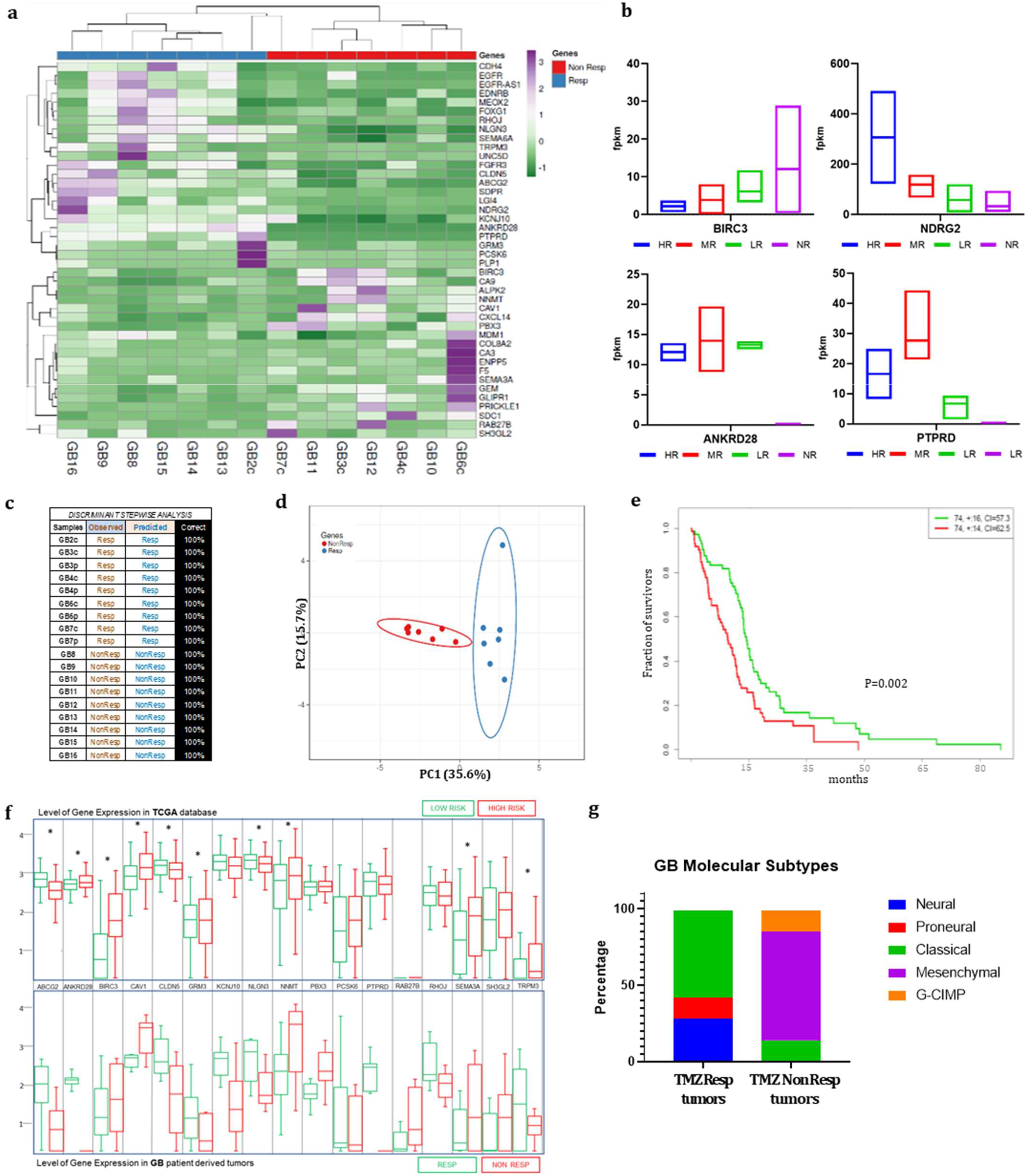
Gene expression profiling of GB-EXPs. (a) Heatmap of 42 genes from RNAseq data differentially expressed (DE) between Resp (n=7) and Non-Resp (n=7). (b) Expression levels of 4 genes subdivided in HR, MR, LR and NR. (c) Stepwise discriminant analysis identifies a combination of 17 DE genes that discriminates the Resp (n=9) and Non-Resp (n=9) cases with 100% accuracy (peripheral portions were included). (d) PCA showing RNAseq data of 17 DE genes in 7 Resp and 7 Non-Resp (tumor peripheral portions were excluded (e) The 17 gene panel Kaplan-Meier plot that identifies a Low-Risk and a High-Risk survival group. Concordance Index = 64.34, Log−Rank Equal Curves p=0.001853, R^2=0.192/0.998 Risk Groups Hazard Ratio = 1.79 (conf. int. 1.23 ∼ 2.6), p=0.002172. (f) Expression levels of the 17 discriminant genes in TCGA database (in the top) and in GB tumor (9 Resp and 9 Non-Resp. in the bottom). (g) Molecular subtypes of GB samples in Resp (n=9) and Non-Resp (n=9) groups.

Finally, on the 42 significant genes, we performed a stepwise discriminant analysis that enabled the identification of a 17 gene signature (Table 6S), which could discriminate the Resp and Non-Resp tumor samples with 100% accuracy (Figure 5c,d). To investigate the potential disease course of patients whose tumors we had predicted to respond to TMZ, we queried the TCGA database of hundreds of clinically and molecularly characterized GB patients. Therefore, the 17 gene profile was used in the “SurvExpress Biomarker Validation of Cancer Gene Expression” tool, to interrogate an extended TCGA-derived GB patient population (n=146). non-parametric statistics, used to estimate the survival function from lifetime data of the 146 cases, produced a Kaplan-Meier plot identifying a low-and high-risk survival group, as shown in Figure 4e,f, whose molecular profile corresponded to the Resp and Non-Resp tumor groups, respectively (Figure 5f). As shown in Figure5e, it is noteworthy that the Resp gene expression profile was associated exclusively with patients that belonged to the group of the longest survivors (>50 months).

To establish the type of GB molecular subtype to which our samples belong, we exploited a TCGA 490 gene expression profile deposited by the Anderson Cancer Center and created a classifier that allowed the assignment of each sample to one of the mesenchymal, classical, proneural, neural, and G-CIMP categories. As shown in Figure 5g, the Non-Resp group was mostly composed of samples of the mesenchymal type, which represents the most aggressive subtype^37^ (Table 7S).

#### Whole Exome Analysis (WEA)

WEA was evaluated in 16 GB tumors divided into nine Non-Resp and seven Resp cases. The mutational landscape of Resp vs. Non-Resp is shown in Figure 6a,b. In Non-Resp samples, the mutational load was higher for any type of variant than in Resp cases (Figure 6a), as widely described in more aggressive and less TMZ-responsive tumor phenotypes^38^. The distribution of base substitutions revealed a prevalence of T>G and C>T transversions in Non-Resp versus Resp tumors (Fig. 6b).

**Figure 6.**
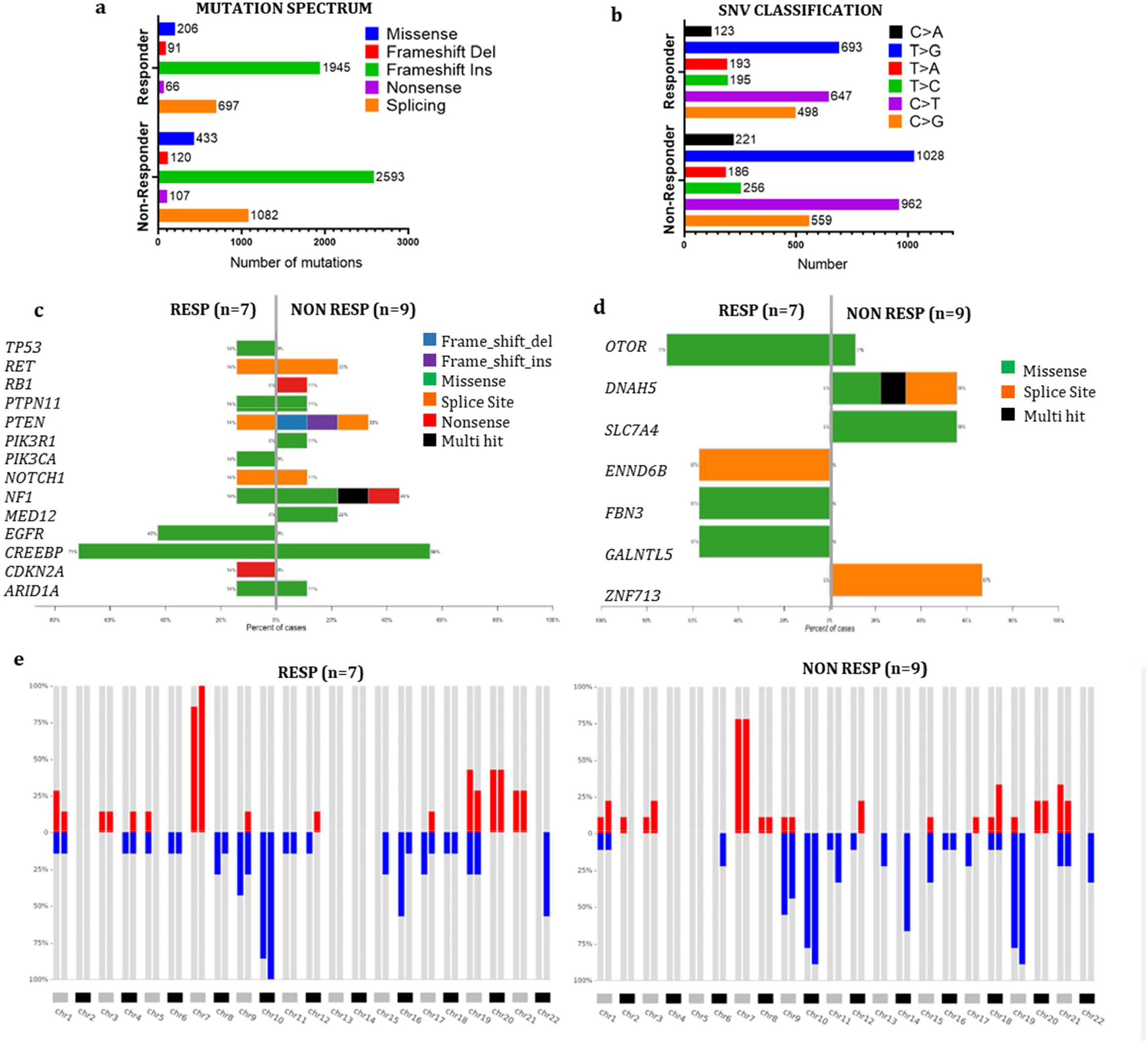
Mutational profiling of GB-EXPs. (a,b) The mutation landscape of the GB cohort (n=16). Counts of each variant classification (a) and counts of each single-nucleotide variant (SNV) classification (b). (c) Co-bar plot of the most frequent gene mutations in GB. (d) Co-bar plot of the genes significantly discriminating Non-Resp and Resp groups. (e) CNApp frequencies for the p and q arms of each chromosome in Resp and Non-Resp groups.

We analyzed mutations in genes known to be altered in IDH1-WT GB by referring to the My Cancer Genome-MCG (mycancergenome.org database). We found 14 mutated genes in the 16 GB cases, for a total of 28 different variants (Figure 6c, Supplementary Figure 5, Supplementary Table 8). The analysis confirmed a higher mutational burden in the Non-Resp group with more deleterious alterations in the PTEN, RB1, and NF1 genes, which are well-known tumor suppressor genes in GB, indicating a more aggressive phenotype in accordance with our previous results ^39^.

Using Maftools forestPlot, we identified seven genes that significantly distinguished between the two groups of tumors (Figure 5d, Supplementary Figure 6, and Supplementary Table 9). Each variant is shown in detail in Supplementary Table 9. Two variants of FBN3, FBN3^E492K^ in GB14 and FBN3^R2688Q^ in GB16, have already been described and annotated in COSMIC with the IDs COSM9337963 and COSM3541504, respectively. The impact on protein function and thus clinical significance has not yet been annotated for most of these variants, resulting in “unknown significance” for the Varsome classification. Only the two splicing variants in DNAH5 in samples GB6c and in DENND6B in sample GB13 were predicted to be pathogenic. Three genes (ZNF713, DNAH5, and SLC7A4) were mutated only in the Non-Resp group (67% (6/9), 56% (5/9), and 56% (5/9), respectively). GALNTL5, FBN3, and DENND6B were shared only by the Resp group at a frequency of 56%. The OTOR gene was common between the groups of tumors, but with a higher mutation rate in Resp cases. To the best of our knowledge, none of the identified genes have been associated with glioblastoma.

CNApp^40^ analysis was used to analyze chromosomal abnormalities in the 16 GB samples (Figure 6e). More than 50% of the samples in both Resp and Non-Resp groups had alterations in chromosomes 7, 9, and 10, which are characteristic of IDH-WT GBs^41^. Non-Resp tumors showed loss of chromosomes 13 and 14, unlike in the Resp group (Figure 6e). Chromosome 14 has been described in GB patients as a carrier of several tumor suppressor genes^42^. Higher gain of chromosomes 19 and 20 was observed in the Resp group than in the Non-Resp group. Amplification of chromosome 19 has been identified as a favorable prognostic marker for GB^43^. These results are indicative of a differential genetic background associated with the FLIM-based TMZ response assessment, further confirming the validity of our approach.

## Discussion

Glioblastoma is the most aggressive malignant tumor of the central nervous system and has a highly unfavorable prognosis. Despite the hardworking search for therapeutic strategies to reverse the highly unfavorable prognoses of GB patients, maximal surgery resection, standard temozolomide chemotherapy (TMZ) and radiotherapy (RT), while not resolutive, currently remain the best treatment option, unchanged since 2005 ^3^. This therapeutic standstill in the GB field is because, novel experimental approaches have shown limited success in improving patient survival^44^. To date, preclinical ex vivo drug testing approaches have failed mainly because they do not reflect the complexity of each individual glioblastoma cellular organization and composition ^5^. In this study, we have developed a novel approach to test the response to an anticancer treatment in patient-derived glioblastoma 3D organoids that we term GB-EXPs. The uniqueness of our approach is to avoid keeping the organoids in culture for extended time to prevent it from undergoing the usual genetic and morphological evolution and divergence from the tumor of origin. Many other in vitro glioblastoma organoid models ^6 8^ consist of several weeks of culturing, wasting the patients’ precious time who in the meantime progresses in their short-term fatal clinical course. This long culture period is also due to the fact that conventional in vitro drug testing assays require very long application settings and readout times with various biological, molecular, genetic, and chemical assays ^45^, that inevitably lead to prolongation of in vitro tumor growth, which in the long term alters an already formed 3D tumor structure, such as a GB-EXP example.

Here we applied a metabolic imaging method that exploits the intrinsic auto-fluorescence molecular properties of NAD(P)H, a metabolic enzyme cofactor, that is associated with the metabolic state of the tissue. Studying intracellular metabolic shifts allows a precocious assessement of cellular response to treatment because anticipates any actual cellular behavior ^11^. In several recent studies, measurement of cancer cell metabolism by live imaging using intrinsic fluorescence from metabolic enzymatic cofactors such as NAD(P)H has shown promise as a sensitive non-invasive method for the early prediction of drug response^10 46^. Therefore, FLIM NAD(P)H, that does not require any type of staining, is perfectly suitable for GB-EXPs because it identifies, at an early stage of culturing, metabolic changes rapidly and non-invasively on the biological material used, leaving it vital ^10^ without interfering with its internal structure ^47 27^. Our approach, therefore, overcomes the limitations of other in vivo drug testing tools since it allows to give a response to treatment after 72hr at the latest from initial treatment and within one week after surgery, allowing the tumor to remain viable and not diverge excessively from how it was structured in vivo in the patient, as shown in Figure 1k,l.

Skala et al. in 2017 implemented for the first time FLIM-based metabolic imaging as an ex-vivo drug testing tool on breast cancer organoids leveraging on NAD(P)H and FAD as metabolic intracellular biomarkers^27^. Here we applied the same approach for the first time on glioblastoma organoids and unlike these authors we analyzed the FLIM measurements using the phasor data analysis approach, a mathematical method that is more suitable for a complex in vitro system such as cancer organoids ^17^. The phasor analysis approach allowed, through the segmentation of the mean distribution curves of the free/bound NAD(P)H fractions operated by the FLIM computational analysis system, the analysis of each tumor in its part and the evaluation of the different internal metabolic states. In such a way, we could calculate the percentage of drug response for each sample and refine the stratification of the tested tumor samples.

We used TMZ to validate the whole procedure. TMZ treatment was the first and only drug to which each GB tumor sample was exposed after first diagnosis. Each patient-derived tumor was classified as a TMZ Resp or Non-Resp sample. We could successfully corroborate our FLIM-based results using conventional drug testing methods and genomic and transcriptomic characterization analyses. In this validation process, a specific molecular status significantly distinguished the two TMZ Resp and Non-Resp tumor populations, stratified phenotypically solely by the NAD(P)H-FLIM based readouts. This distinction at the molecular level strengthens the accuracy of this approach. In fact, between the two groups of patient-derived tumors, which share many common characteristics of GB, we have been able to highlight, quite strikingly, markers already described in the literature, but also new potential targets associated with the response to TMZ, such as ANKRD28, PTPRD, ZNF713, DNAH5, SLC7A4, GALNTL5, FBN3, DENND6B, and OTOR. A unique new 17-gene expression signature significantly discriminating the Resp and Non-Resp groups emerged. Since our tumor cases derived from patients for whom the clinical course was not yet available, we could not have a clinical confirmation of what was predicted in vitro. Therefore, in the meantime, we decided to test the tumor molecular profiles on a series of 150 patients of the glioblastoma TCGA dataset completely clinically characterized. This investigation revealed that the molecular profile that characterized our TMZ responsive and non responsive tumor populations were highly significantly associated with the long-surviving and short surviving groups of the TCGA dataset, as we would predict. Further studies are required to investigate the potential role of the 17 genes signature in glioblastoma.

Furthermore although MGMT methylation status is accepted as the only molecular prognostic biomarker for predicting patient response to TMZ treatment, inconsistencies do occur and currently challenge the efficacy of this biomarker in clinical practice, raising the question of its value^48^. Here the number of methylated tumors for MGMT was higher in the responsive group than in the non-responsive group, as expected. However, following our procedure, some non-methylated tumor cases turned out to be nevertheless responsive to TMZ, suggesting that our approach could be synergistic with the classical MGMT methylation biomarker.

This study is the first to apply FLIM-based metabolic imaging to in vitro vital patient-derived GB tumors to perform an ex vivo treatment tumor response assessment early before losing the parental tumor architecture. The lack of predictive biomarkers is a major limitation in GB clinical oncology, and today, clinicians urgently need step-changing informative tools to support their decision-making therapy approaches. A method to predict a patient’s tumor-specific drug response before the onset of therapy can be useful for managing patients with GB. This innovative approach in the field of glioblastoma can be transformative for the clinical management of the patients with glioblastoma, especially at the time of disease progression when the guidelines are less stringent and when the patient can receive more therapeutic options. The performance of such a functional precision medicine approach can provide additional information regarding a patient’s tumor vulnerabilities. These functional approaches open the door to new discoveries and generate further knowledge of the disease, which in turn will increase the likelihood of producing useful therapeutic solutions ^49^. It is important to point out that the full promise of precision medicine in oncology is yet to be realized, as more individuals may benefit from functional approaches. In the glioblastoma clinical field, we envision an increasing implementation of functional precision medicine protocols and a transition from therapeutic approaches followed by watchful waiting to informed decisions based on specific patient-derived GB tumor treatment-response predictions. This approach can also be used to test new FDA-approved anti-cancer drugs in vitro directly on the tumor, could also be seen as a springboard for new drugs that need to be transferred to more advanced stages of clinical trials and be implemented for other cancers.

## Acknowledgements

We would like to thank Prof. Kevin Janes at the Department of Biomedical Engineering, University of Virginia, USA, for kindly providing us with OrganoSeg Software. We would also like to express our special thanks to Mr. Raffaele Cotugno for his kind donations to this project.

## Notes

Funding Statement: The study was supported by FPSgrant2018 from the Fondazione Pisana per la Scienza

### Competing Interest Statement

The authors have declared no competing interest.

