## Supplementary Materials for "Metabolic-imaging of human glioblastoma live tumors: a new precision-medicine approach to predict tumor treatment response early"

### SUPPLEMENTARY MATERIAL AND METHODS

#### Human glioblastoma tissue collection

Each tumor sample was washed with Dulbecco's phosphate-buffered saline (DPBS) in a sterile dish and portioned with a scalpel into ~ 0.5-1 $mm^2$ pieces under a biological hood. We have developed a procedure to minimize the tampering of samples. In an effort to minimize variability due to sampling, we pooled 2-4 pieces of parental tumor tissue into one sample for the subsequent analysis: A summary chart of the specific analyses performed for each tumor sample, including FLIM analysis, H&E staining, drug treatment, radiation treatment, size analysis, Ki67 expression analysis, MGMT methylation testing, whole transcriptome analysis (WTA), whole exome analysis (WEA) is reported in Table 3S. The surgery tumor tissue if not immediately processed, was viable frozen at -140°C in 90 % fetal bovine serum (FBS) and 1% dimethyl sulfoxide (DMSO). The tumor piece for histological analysis was immediately fixed in 10% formalin and embedded in paraffin and the portions for the remaining analyses were stored at -80°C. Notably, we cultured GB-EXPs directly right after resection but also after short time storage at -140°C, confirming that viable cultures can be grown either from fresh and flash-frozen DMSO supplemented media, thus facilitating the whole procedure [1]. Samples underwent several analyses as shown in Table 3S.

#### Glioblastoma Cell Lines

T98G and U87 GBM cell lines were obtained from American Type Culture Collection (ATCC, Rockville, MD). T98G and U87 were grown as monolayers in Dulbecco's Modified Eagle Medium (DMEM) low glucose and high glucose respectively without red phenol, supplemented with 10% FBS and 1% Penicillin-Streptomycin. For FLIM experiments, cells were grown in 35 mm Nunc Glass Bottom Dishes (Thermo Fisher Scientific).

The identity of each commercial cell line was verified with Gene Print 10 System (Promega), through STR analysis of ten different loci, including all ASN-0002-2011 loci (TH01, TPOX, vWA, CSF1PO, D16S539, D7S820, D13S317, D5S818) plus Amelogenin and D21S11. Detection of amplified fragments was performed using the Applied Biosystems 3500 Genetic Analyzer (Applied Biosystems) and analyzed using GeneMapperID software (Applied Biosystems). STR profiles of the analyzed cell lines were compared to STRs available in ATCC online databases.

#### Confocal Lifetime Imaging

Images were taken with Olympus Fluoview 3000 confocal microscope, equipped with four laser lines (405/488/561/640 nm), 2 hybrid detectors and 2 standard detectors (Olympus, FV31-

HSD and FV31-SD), using a quadriband 405/488/561/640 nm dichroic mirror (Chroma) and an UPLXAPO20X (20X magnification, N.A.=0.80) for brightfield acquisition or UPLXAPO60XO (60X magnification, N:A.=1.42) oil immersion objective for FLIM. Confocal pinhole diameter was set to 1 Airy. Fluorescence lifetime imaging was performed using MultiHarp 150 (Picoquant) time-correlated single-photon counting (TCSPC) unit, controlled through SymPhoTime64 software (PicoQuant), using a 405 nm LDH-P-C-375B (Picoquant) excitation laser for NAD(P)H [23]. Fluorescence was collected with two PMA hybrid detectors (Picoquant) using a dichroic filter (510 nm), and bandpass filters 440/40 for NAD(P)H. Laser pulse frequency was set to 40 MHz, pixel dwell time was set to 10 us, and 240 cycles of acquisition were performed for each field. Images sizes were of 512X512 pixels. Temporal resolution was 80 ps.

#### Cell viability assay

Cell viability was determined using the WST1 assay (Clontech Laboratories, Mountain View, CA, USA). A total of 5000 cells per well were seeded in a 96-well plate format. At the time of seeding (T0) and after 24 h (T1), 48 h (T2) and 72 h (T3), the WST1 reagent was added and incubated for a further 60 min before reading the plate. Each assay was conducted in triplicate. The quantity of formazan dye is directly related to the number of metabolically active cells and was quantified by measuring the absorbance at 450 nm in a multiwell plate reader (Tecan, Mannedorf, Switzerland). OD values at 24 h (T1), 48 h (T2) and 72 h (T3) were normalized to T0.

#### Nucleic Acids Isolation

DNA extraction was performed from parental tissue sample, stored at -80 °C, using Maxwell 16 Tissue LEV DNA Purification Kit (Promega, Madison, WI, USA) according to manufacturer's protocol. RNA was extracted from parental tumor samples, GBM cells, explants and spheroids in suspension using Maxwell 16 LEV Simply RNA Tissue Kit (Promega, Madison, WI, USA), according to manufacturer's protocol. DNA and RNA concentrations were determined using the Qubit Fluorometer (Life Technologies, Carlsbad, CA) and the quality was tested using the Agilent 2200 Tapestation (Agilent Technologies, Santa Clara, CA) system.

#### Ki67 Expression analysis

Complementary DNA (cDNA) was prepared from 2 ng of total RNA via reverse transcription using iScript cDNA Synthesis Kit (Bio-Rad) in a final volume of 20 µl according to manufacturer's protocol.

To study Ki67 expression in cell lines we performed semiquantitative real-time PCR in a 10 µL reaction mixture consisting of 5 µL of SsoAdvanced Universal SYBR Green supermix (Bio-Rad), 1 µL of primer Assay (Bio-Rad), 2.0 µL of cDNA, and 2 µL of nuclease-free water. We used Human Mki67 PrimePCR SYBR Green Assay (Bio-Rad) for Ki67 and Human ACTB PrimePCR™ SYBR® Green Assay (Bio-Rad), for B-actin housekeeping gene. Each PCR amplification was performed using CFX96 Touch Deep Well PCR system (Biorad), at the following conditions: initial template denaturation at 98 °C for 30 s, followed by 40 cycles of denaturation at 98 °C for 15 s and annealing at 60 °C for 30 seconds. Each sample was assessed in triplicate, and positive/negative controls were included in parallel for each reaction. Amplification was followed by melting curve analysis to assess PCR product specificity. The data were analyzed using the $2^{-\Delta\Delta CT}$ method of relative quantification[4].
For Ki67 expression analysis in explant cultures and spheroids we used Digital Droplet PCR method (DDPCR). The BioRad QX200 Droplet Digital PCR system (Bio-Rad Laboratories, Inc.) was used to perform DDPCR. In each DDPCR reaction 10 ul of 2x digital PCR supermix for probes (No dUTP) (Bio-Rad), 1 ?l of each gene expression assay, 2 ?l of cDNA and 6 ?l of RNAse free water were used. MKI67 Human ddPCR Gene Expression Assay (Bio-Rad), having a FAM-labeled probe, was used to detect Ki67 transcripts, while BACT Human ddPCR Gene Expression Assay (Bio-Rad), having an HEX-labeled probe, was used as housekeeping to amplify B-actin transcript. Each sample was analyzed in triplicate. PCR was performed using C1000 Touch Thermal Cycler (Bio-Rad) with the following conditions: 95°C for 10 min; followed by 40 cycles of 94°C for 15 sec and 58°C for 60 sec; and a final extension at 98°C for 10 min. After thermocycling, the 96-well plate was put in the plate holder and read in the QX200 Droplet Digital PCR system, and based on positive droplets and according to the Poisson distribution, the absolute copy number of Ki67 and of B-actin was calculated using QuantaSoft™ analysis software (version 1.7.4.0917; Bio-Rad Laboratories, Inc.). Ki67 expression levels are reported as Ki67/B-actin ratios.

**Live/Dead Assay**

The assay was performed using Molecular Probes LIVE/ DEAD viability/cytotoxicity kit (Molecular Probes). After incubation explants were visualized under Olympus Fluoview 3000 microscope. 515 nm for calcein and 635 nm for EtD-1 optical filters were used to image the explants at a 20X magnification.

**Histology and staining**

Tissues and explants, both in matrigel and in suspension, were fixed for 24 h, in 10% neutral-buffered formalin (Sigma-Aldrich) at room temperature, processed through a graded ethanol series followed by xylene, and embedded in paraffin. Paraffin-embedded samples sections (5 µm) were stained with hematoxylin (Diapath C0303) for 40 s and with eosin (Diapath C0353) for 30 s. For immunohistological staining, paraffin slides were deparaffinized and subjected to antigen retrieval using Epitote Retrieval Solution (ph=8) (Leica Microsystems RE 716 CE). Samples were incubated with Ki67 monoclonal (SP6) (Thermo Fischer, MA5-14520) primary antibody using 1:50 dilution for 1 h at RT. Detection of bound antibody was accomplished with the Rabbit Specific HRP/DAP Detection IHC Kit (Abcam, ab64261) Immunohistological and H&E pictures were taken with microscope (CARL ZEISS Axio Observer Z1FLMot) after mounting with mounting medium (Fisher Scientific, 7 Miami, FL). For immunofluorescence, the paraffin blocks were sliced into 5 μM thick sections, deparaffinized with xylene (Fisher Scientific, Waltham, MA, USA), and rehydrated with decreasing concentrations of ethanol in water. For explants in matrigel, the medium was removed and samples were washed twice with DPBS solution. The antigen unmasking was achieved with Epitote Retrieval Solution (ph=8) (Leica Microsystems RE 716 CE) in a microwave. CD105 polyclonal (Thermo Fisher, PA5-94980), CD3 monoclonal (F7.2.38) (Thermo Fisher, MA5-12577), Sox2 polyclonal (Thermo Fisher, 48-1400), GFAP monoclonal (ASTRO6) (Thermo Fisher, MA5-12023), GFAP polyclonal (Abcam, ab7260), Iba1 polyclonal (Wako 091-19741) primary antibodies were then applied at dilutions of 1:400, 1:20, 1:100, 1:100, 1:250, 1:1000, respectively overnight at 4 °C. Goat anti-rabbit Alexa Fluor 488 (Thermo A11034) and Goat anti-mouse Alexa Fluor 568 were diluted 1:500 and incubated for 1h. Cells were counterstained with DAPI (Sigma F6057) and visualized using Olympus Fluoview 3000 confocal microscope at a magnification of 60X, as above described.

**Next-Generation Sequencing data analysis**

RNA-Seq reads in FASTQ format were initially inspected using FASTQC program (http://www.bioinformatics.babraham.ac.uk/projects/fastqc/). Reads were mapped against the reference genome (Hg19) by using STAR aligner (version 2.5.3a). Read counts on known human genes were calculated by feature Counts version 1.5.1[5]. Differential expression was performed using two independent tools: edgeR 2.6.12 [6], and Cuffdiff 2[6]. Differentially expressed (DE) genes were selected intersecting lists of significant DE genes with p-value < 0.004 obtained by CuffDiff2 and edgeR. A discriminant stepwise analysis was used to find genes discriminating our two groups: Resp and Non-Resp. JMP 10.0.0 (SAS Institute). The same statistical approach was used for classifying GB molecular subtypes leveraging on the TCGA Anderson Cancer Center. . To compare the prognostic significance of our discriminant gene set in predicting survival in glioblastoma patients, we evaluated it in TCGA Database with SurvExpress platform. Heatmaps and PCA plots were generated using ClustVis[7]. The primary data analysis of exomes was performed by using the SeqMule pipeline[8]. Somatic single-nucleotide variants (SNVs) and indels were identified in tumors against a PoN (Panel of Normal, GATK best practices, https://console.cloud.google.com/storage/browser/gatk-best-practices/somatic-b37) by using Mutect2 [9] variant calling algorithms. Rare variants were obtained excluding somatic variants reported in the non-cancer database gnomAD v3[10] setting minor allele frequency (MAF) of ≥0.01. The frequency and type of mutations were investigated using the R package MAFtools[11]. Copy number was estimated by CNVkit[12]. Copy number variations have been summarized using CNApp[13] with default cutoffs. Comparison data for CNV classifier were downloaded from The Cancer Genome Atlas Glioblastoma Multiforme (TCGA-GB, https://www.cancer.gov/tcga) data collection (hg19 Legacy Database) using the

**Methylation-specific multiplexed ligation-dependent probe amplification (MS-MLPA) for MGMT promoter methylation analysis**

The methylation status of the MGMT promoter was tested using the Salsa MS-MLPA kit (MRC-Holland, Amsterdam, The Netherlands), as described by the manufacturer. The kit contains three probes specific for the MGMT promoter region, including HhaI recognition site: MGMT1 (5670-L5146; 193 bp), MGMT2 (2239-L1261;373 bp), and MGMT3 (5668-L5144; 454 bp). Briefly, after denaturation of the sample, probes were hybridized and then ligated. For half of the sample, ligation was combined with HhaI digestion. HhaI is a methylation-sensitive restriction enzyme that cuts unmethylated GCGC sites. The resultant polymerase chain reaction fragments were separated by capillary gel electrophoresis (3500 Genetic Analyzer; Thermo Fisher). The methylation status was quantified using GeneMapper software (version 1.5; Soft Genetics, State College, PA). To evaluate the methylation status, the "methylation ratio" was calculated by dividing each normalized peak value of the HhaI enzyme digested sample by that of the corresponding undigested sample. This value corresponds to the percentage of methylated sequences.

### SUPPLEMENTARY DATA

### SUPPLEMENTARY FIGURES

**Supplementary Figure S1:** (a) The phasor plot representation creates a heatmap located in the center of the plot inside the universal circle by clustering the pixels of the image that have similar lifetimes. Positioning cursors on the universal circle in the free- and protein bound-NAD(P)H position, a metabolic trajectory is created (black arrow) and according to the protocol described by Ranjiit et al. [14] we can extrapolate from phasor plot the NAD(P)H fractional distribution curves (see materials and methods for pixel phasor plotting procedure). (b) Positioning the two cursors at the extremities of the phasor plot, the metabolic trajectory from NAD(P)H in the bound state (red/magenta) to NAD(P)H in the free state (green/white) was represented in a phasor map as reported by the color-bar on the side.

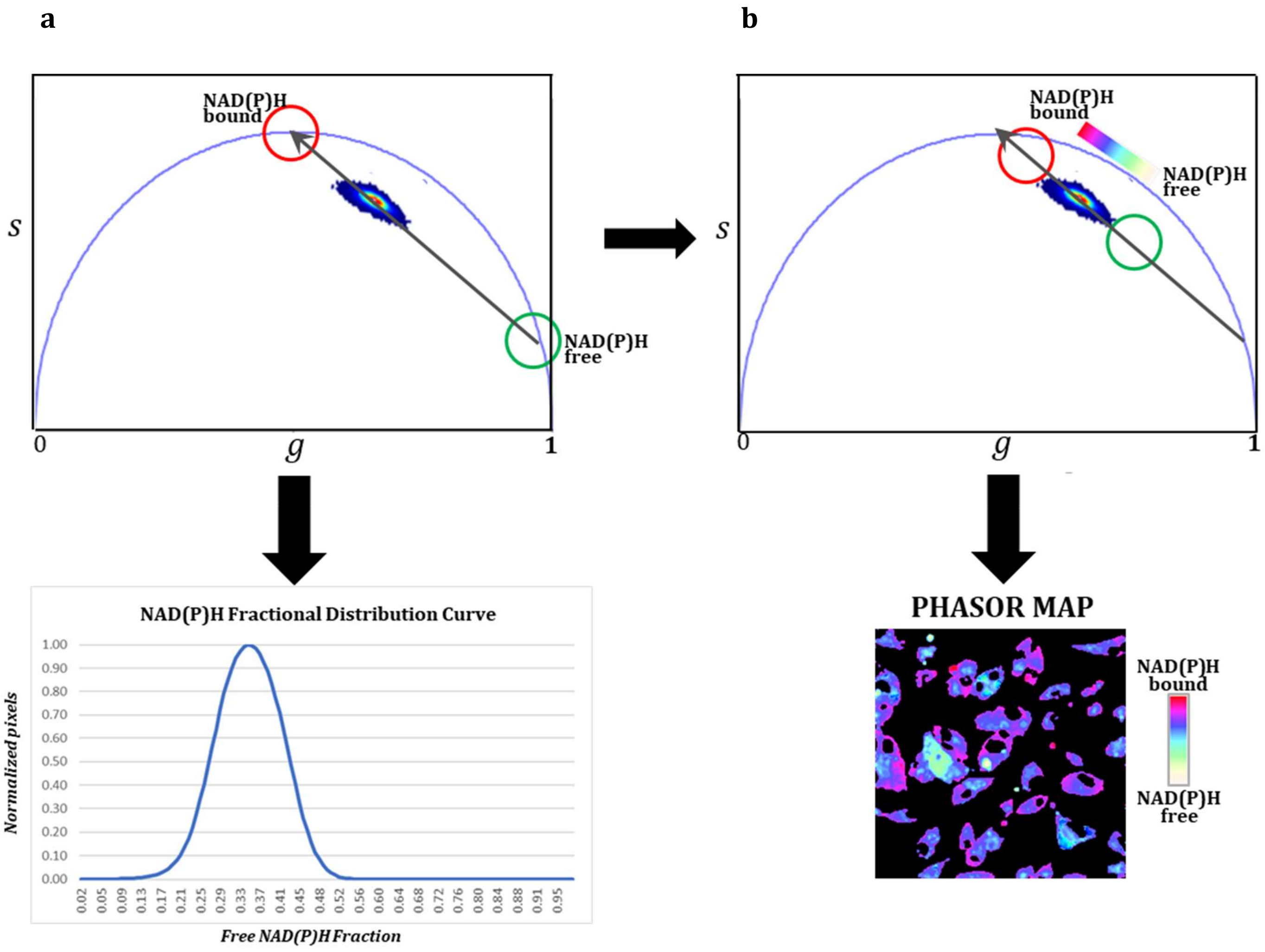

**Supplementary Figure S2**: Histological pictures of IDH1/2 WT surgery GB tissues. Hematoxylin and Eosin staining is shown. Scale bar, 125 µm.

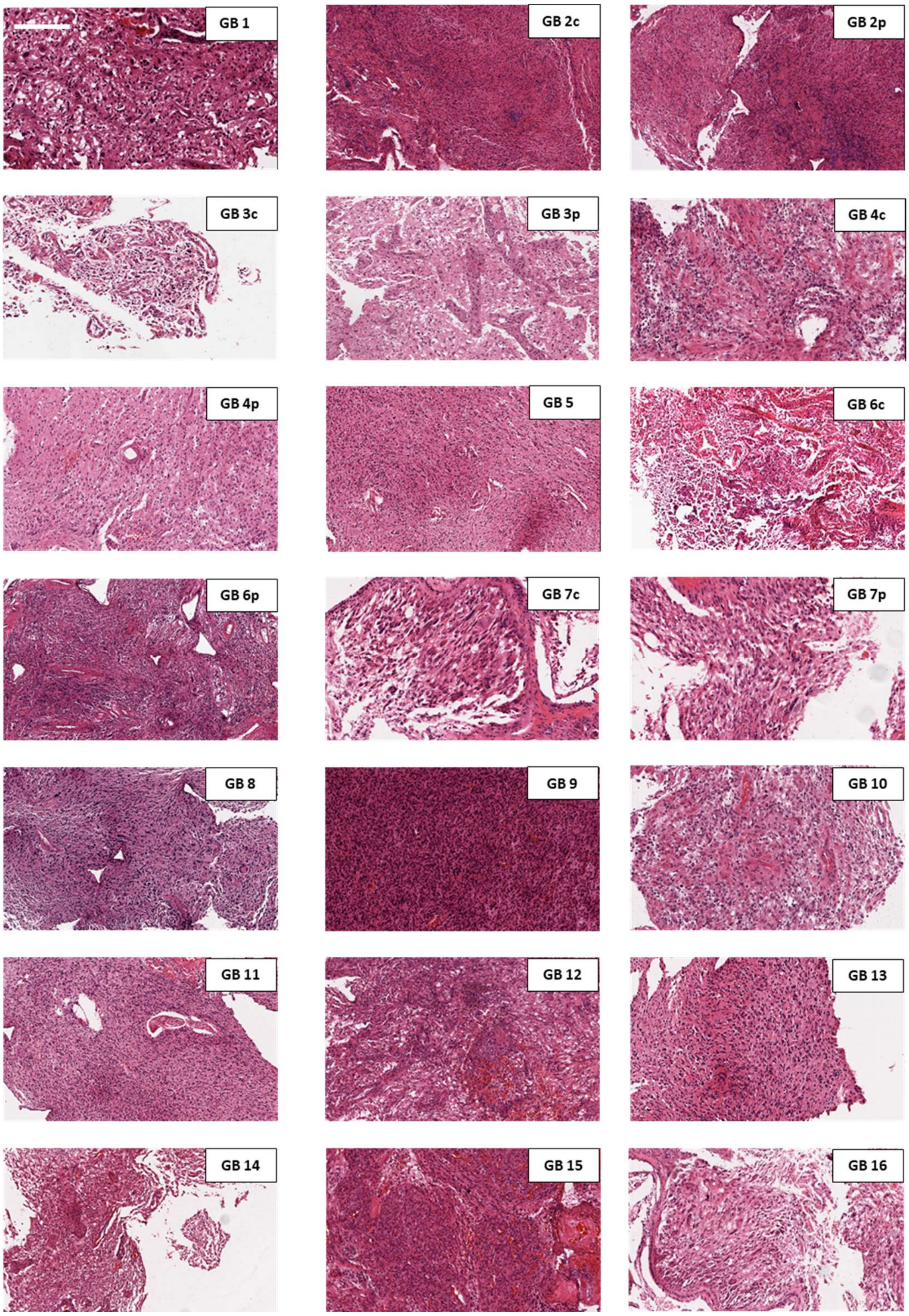

**Supplementary Figure S3**: Representative images of GB-EXPs (a,b) and details (c-f) showing their vitality at 2 weeks after culturing. Scale bar, 40 µm.

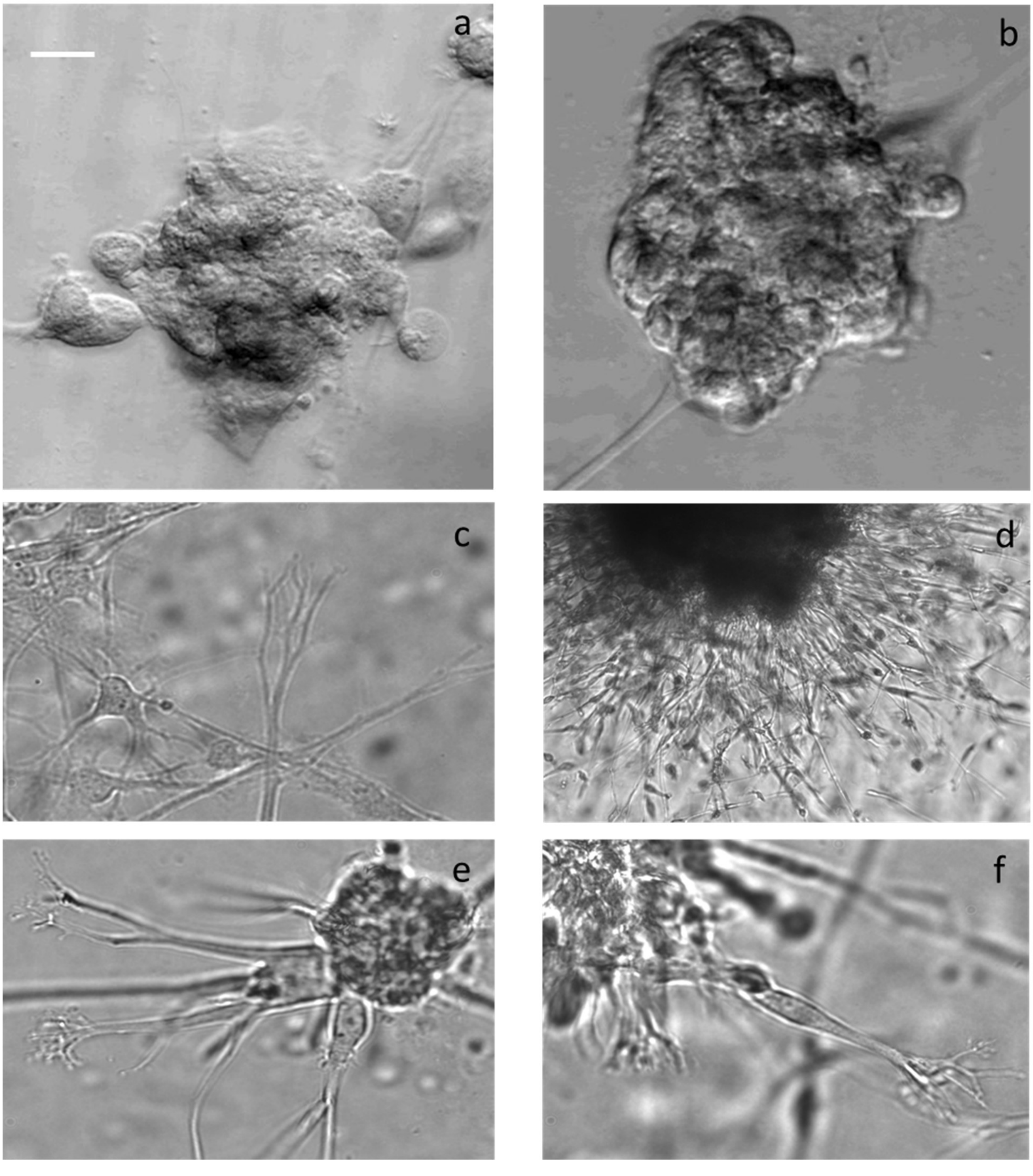

**Supplementary Figure S4**: Comparison of Final %DR in tumor core and peripheral-derived GB –EXPs of 5 GB samples.

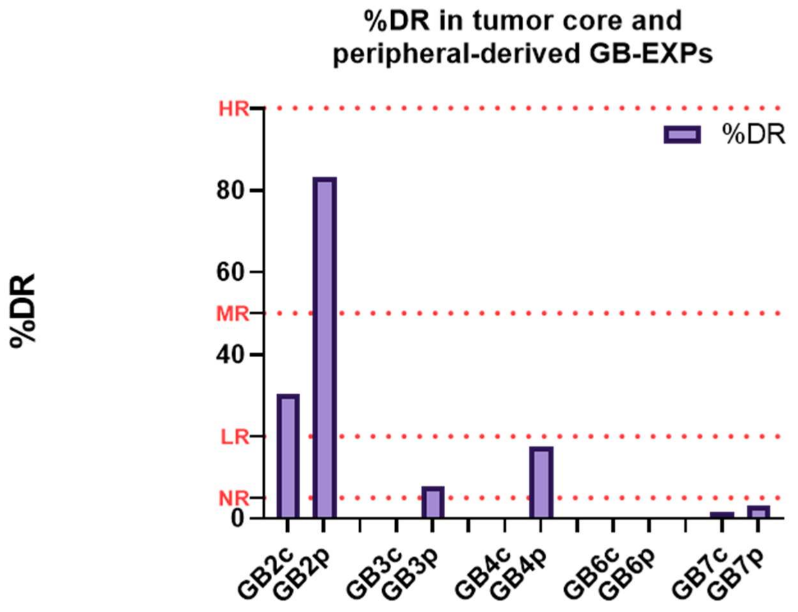


According to %DR: NR<5%; 5≤LR<20; 20≤MR<50; HR≥50

**Abbreviations**: NR= Non-Resp; LR= Low Resp; MR=Medium Resp; HR= High Resp

**Supplementary Figure S5**: Oncoplot of the distribution of mutations found in our samples in the most frequently mutated genes in GB IDH1 WT. Each column represents one sample and each row a different gene. Colored squares show mutated genes, while empty (gray) squares show no mutated genes. The different types of mutations are colored according to the type of variant: orange, splice site mutation; blue, frameshift deletion; green, missense mutation; red, nonsense mutation; and black, multi-hit mutation. Genes annotated as “multi-hit” have more than one type of mutation in the same region.

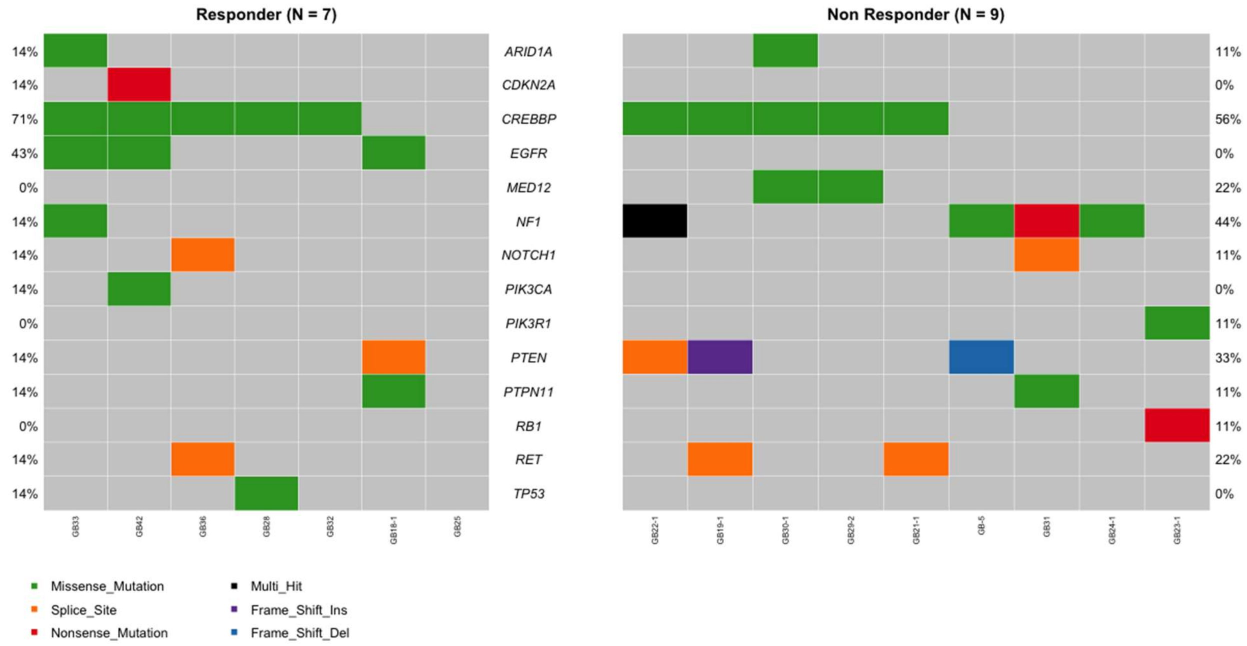

**Supplementary Figure S6**: Oncoplot of statistically significant genes that differentiate the two groups Non-responder and Responder. Each column represents one sample and each row a different gene. Colored squares show mutated genes, while empty (gray) squares show no mutated genes. The different types of mutations are colored according to the type of variant: orange, splice site mutation; blue, frameshift deletion; green, missense mutation; red, nonsense mutation; and black, multi-hit mutation. Genes annotated as “multi-hit” have more than one type of mutation in the same region.

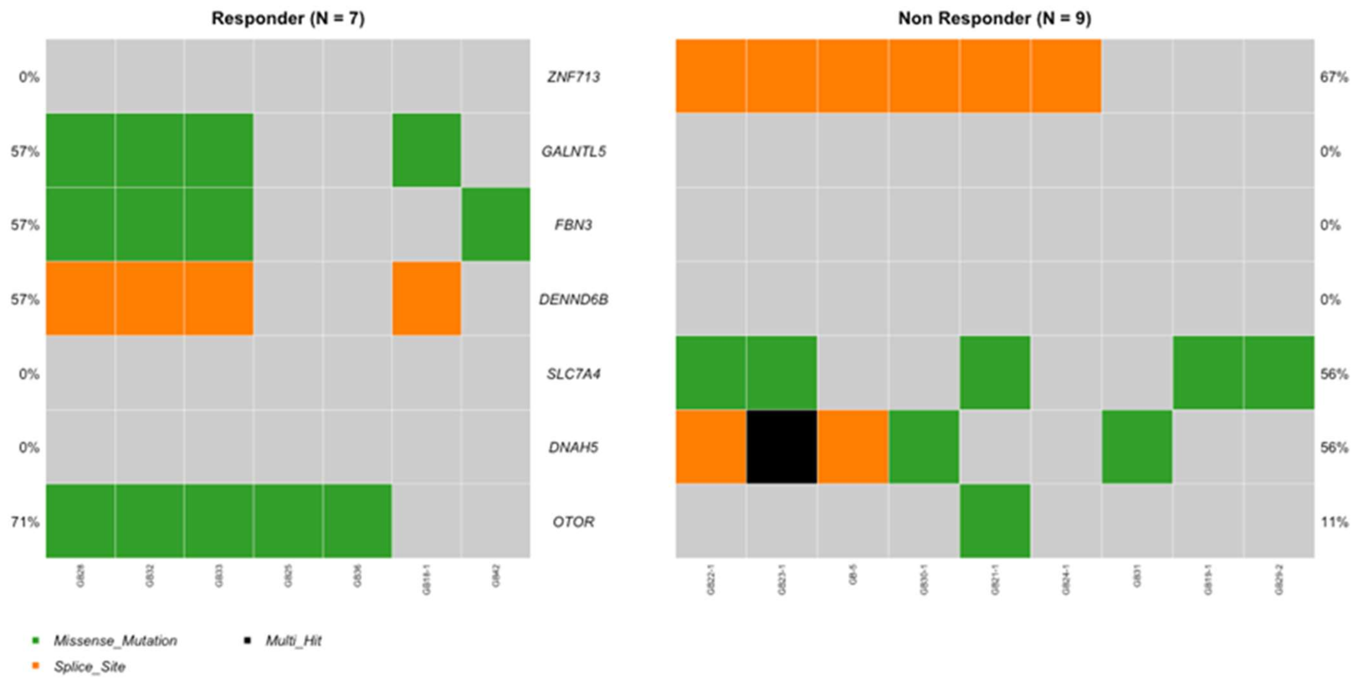

### SUPPLEMENTARY TABLES

**Table S1** Patients' clinical and demographic data

| Cases | Periphery(p)/ core (c) | Age | Sex | Primary/Reccurent | Brain Location | IDH1/IDH2 | MGMT methylation status* performed on surgery GB tissues |
|---|---|---|---|---|---|---|---|
| **GB1** | / | 62 | M | Primary GB | right parietal lobe | WT | NM (5%) |
| **GB2** | c | 61 | M | Primary GB | left temporal lobe | WT | NM (6%) |
| | p | | | | | | / |
| **GB3** | c | 69 | F | Primary GB | rolandic frontal lobe | WT | NM (2%) |
| | p | | | | | | M (33%) |
| **GB4** | c | 60 | F | Primary GB | left temporal lobe | WT | NM (4%) |
| | p | | | | | | M (36%) |
| **GB5** | / | 73 | M | Primary GB | temporo-parietal lobe | WT | M (18%) |
| **GB6** | c | 69 | F | Primary GB | left occipital lobe | WT | NM (2%) |
| | p | | | | | | M (14%) |
| **GB7** | c | 74 | M | Primary GB | right temporal lobe | WT | NM (4%) |
| | p | | | | | | M (38%) |
| **GB8** | / | 57 | M | Primary GB | left frontal-parietal lobe | WT | M (23%) |
| **GB9** | / | 30 | M | Primary GB | parietal lobe | WT | NM (3%) |
| **GB10** | / | 41 | F | Primary GB | right temporo-occipital lobe | WT | M (21%) |
| **GB11** | / | 74 | F | Primary GB | right frontal lobe | WT | NM (7%) |
| **GB12** | / | 47 | M | Primary GB | right temporal lobe | WT | NM (2%) |
| **GB13** | / | 46 | F | Primary GB | parietal lobe | WT | M (11%) |
| **GB14** | / | 71 | M | Primary GB | frontal lobe | WT | M (16%) |
| **GB15** | / | 65 | M | Primary GB | right frontal lobe | WT | M (23%) |
| **GB16** | / | 80 | M | Primary GB | right temporal insular lobe | WT | M (14%) |

_* methylation threshold >7%_

Light blue cells show GB samples MGMT methylation status results different in core and periphery portion of the same GB tumor tissues.

**Abbreviations:** WT=wild-type; c=core; p=periphery; GB=glioblastoma; M=male; F=female

**Table S2** Pathological diagnosis and therapy administrated for all GB patients.

| GB-cases | Pathology Report | Therapy Administered |
|---|---|---|
| GB1 | Glioblastoma (grade IV WHO; GFAP+, p53 normal pattern of expression, MKI67-1 23%, codeletion 1p-19q) | Soldesam, Mannitol 18%, Antra, Cefazoline |
| GB2C | Glioblastoma (grade IV WHO; GFAP+, p53-/+, MKI67-1 30%) | Soldesam, Mannitol 18%, Antra, Cefazoline |
| GB2p | Glioblastoma (grade IV WHO; GFAP+, p53-/+, MIB-1 30%) | Soldesam, Mannitol 18%, Antra, Cefazoline |
| GB3c | Recurrent glioblastoma (Grade IV WHO;GFAP+, p53+, MKI67- 45%) | Soldesam, Lvofloxacine, Antra, Cefazoline |
| GB3p | Recurrent glioblastoma (Grade IV WHO;GFAP+, p53+, MKI67- 45%) | Soldesam, Lvofloxacine, Antra, Cefazoline |
| GB4c | Glioblastoma (Grade IV WHO; GFAP-/+, p53+/-, MKI67-1 40%) | Soldesam, Mannitol 18%, Antra, Cefazoline |
| GB4p | Glioblastoma (Grade IV WHO; GFAP-/+, p53+/-, MKI67-1 40%) | Soldesam, Mannitol 18%, Antra, Cefazoline |
| GB5 | Glioblastoma (Grade IV WHO; GFAP+, p53+,MKI67-1 35%) | Soldesam, Antra, Cefazoline |
| GB6c | Glioblastoma (Grade IV WHO; GFAP+, EGFR+, p53-, MKI67-1 35%) | Soldesam, Mannitol 18%, Antra, Cefazoline |
| GB6p | Glioblastoma (Grade IV WHO; GFAP+, EGFR+, p53-, MKI67-1 35%) | Soldesam, Mannitol 18%, Antra, Cefazoline |
| GB7c | Glioblastoma (Grade IV WHO; GFAP+, p53 normal pattern of expression, MKI67-1 40%) | Soldesam, Antra, Cefazoline |
| GB7p | Glioblastoma (Grade IV WHO; GFAP+, p53 normal pattern of expression, MKI67-1 40%) | Soldesam, Antra, Cefazoline |
| GB8 | Glioblastoma (Grade IV WHO) (GFAP+, MKI67-20%) | Levetiracetam, Desametasone, Lansoprazole |
| GB9 | Glioblastoma (Grade IV WHO) (GFAP+, MKI67-20%) | Levetiracetam, Soldesam, Lansoprazole |
| GB10 | Glioblastoma (Grade IV WHO; GFAP+, p53+/-, MKI67-1 35%) | Soldesam, Mannitol 18%, Antra, Cefazoline |
| GB11 | Glioblastoma (Grade IV WHO; GFAP+, EGFR+, p53+/-, Sox-11+/-, MKI67-1 25%) | Soldesam, Mannitol 18%, Antra, Cefazoline |
| GB12 | Glioblastoma (Grade IV WHO) (GFAP+, MKI67-30%) | Levetiracet,, Dexamethasone, Omeprazole |
| GB13 | Glioblastoma (Grade IV WHO) (GFAP+, MKI67-20%) | Levetiracet, Dexamethasone, Omeprazole |
| GB14 | Glioblastoma (Grade IV WHO) (GFAP+, MKI67-20%) | Omeprazole, Levetiracetam, Lansoprazole |
| GB15 | Glioblastoma (Grade IV WHO) (GFAP+, MKI67-20%) | Levetiracetam, Lansoprazole, Dexamethasone (Mannitol pre-op) |
| GB16 | Glioblastoma (Grade IV WHO) (GFAP+, MKI67-10%) | Phenytoin, Levetiraceta, Soldesam, Lansoprazole, Lacosamide |

**Table S3** Type of analyses performed for each GB case.

| Cases | Explant culture | EXPsTMZ treatment | FLIM on GB samples (24/48/72hr) | OrgaonSeg Size Analysis on GB samples in matrigel (0,1,2 weeks) | Ki67 mRNA expression analysis on GB EXPs (0,2 weeks) | MGMT methylation analysis on GB-EXPs | IDH1/IDH2 on GB-EXPs | GB EXPs H&E | Histology GB EXPs | IHC Pathology Report GB Tissue | WTA on GB EXPs | WEA on GB EXPs | GB EXPs molecular subtype by WTA |
|---|---|---|---|---|---|---|---|---|---|---|---|---|---|
| 2D-T98G | / | x | x | | x | | | | | | | | |
| 2D-U87 | / | x | x | | x | | | | | | | | |
| 3D-T98G | / | x | x | x | x | | | | | | | | |
| 3D-U87 | / | x | x | x | x | | | | | | | | |
| GB1 | x | x | x | x | | x | x | x | x | x | | x | |
| GB2c | x | x | x | x | | x | x | x | x | x | x | x | x |
| GB2p | x | x | x | x | x | | | x | x | x | | | |
| GB3c | x | x | x | x | | x | x | x | x | x | x | x | x |
| GB3p | x | x | x | | | x | x | x | x | x | x | x | x |
| GB4c | x | x | x | x | x | x | x | x | x | x | x | x | x |
| GB4p | x | x | x | x | x | x | x | x | x | x | x | x | x |
| GB5 | x | x | x | x | x | x | x | x | x | x | | x | x |
| GB6c | x | x | x | x | x | x | x | x | x | x | x | x | x |
| GB6p | x | x | x | x | x | x | x | x | x | x | x | x | x |
| GB7c | x | x | x | x | x | x | x | x | x | x | x | x | x |
| GB7p | x | x | x | x | x | x | x | x | x | x | x | x | x |
| GB8 | x | x | x | x | x | x | x | x | x | x | x | x | x |
| GB9 | x | x | x | x | x | x | x | x | x | x | x | x | x |
| GB10 | x | x | x | x | | x | x | x | x | x | x | x | x |
| GB11 | x | x | x | x | x | x | x | x | x | x | x | x | x |
| GB12 | x | x | x | x | x | x | x | x | x | x | x | x | x |
| GB13 | x | x | x | x | x | x | x | x | x | x | x | x | x |
| GB14 | x | x | x | x | x | x | x | x | x | x | x | x | x |
| GB15 | x | x | x | x | x | x | x | x | x | x | x | x | x |
| GB16 | x | x | x | x | | x | x | x | x | x | x | x | x |

**Table S4** Number of images acquired for FLIM analysis and for size analysis for each GB case.

| | Total Number of fields (in cells), spheroids, GB EXPs analysed by FLIM (Ctrl /Treated) | Total Number of GB EXPs images analyzed for Size measurement at 0week, 1week, 2weeks |
|---|---|---|
| 2D-T98G | 10/10 | |
| 2D-U118 | 10/10 | |
| 3D-T98G | 5/5 | 75 |
| 3D-U118 | 5/5 | 33 |
| GB1 | 14/16 | 81 |
| GB2c | 19/19 | 36 |
| GB2p | 19/19 | 54 |
| GB3c | 18/18 | 33 |
| GB3p | 17/18 | 30 |
| GB4c | 18/18 | 48 |
| GB4p | 20/20 | 42 |
| GB5 | 17/15 | 42 |
| GB6c | 18/18 | 41 |
| GB6p | 20/20 | 59 |
| GB7c | 18/18 | 41 |
| GB7p | 24/24 | 65 |
| GB8 | 22/20 | 45 |
| GB9 | 26/28 | 60 |
| GB10 | 30/29 | 69 |
| GB11 | 30/30 | 72 |
| GB12 | 22/22 | 30 |
| GB13 | 30/30 | 15 |
| GB14 | 30/30 | 30 |
| GB15 | 32/33 | 81 |
| GB16 | 28/30 | 72 |

**Table S5** %DR for each patient-derived GB-EXPs (n=21) at 24hr, 48hr, 72hr of TMZ treatment. The final %DR is reported as "Weighted Average". Phasor-FLIM phenotype is shown.

| % DR (HR<5%, 5% ≤MR <20%, 20% ≤LR <50% and NR≥ 50% ) | | | | | |
|---|---|---|---|---|---|
| | **24hr** | **48hr** | **72hr** | **Weighted Average** | **Phasor-FLIM phenotype** |
| **GB1** | 0 | 0 | 0 | 0 | **NR** |
| **GB2c** | 84.51 | 35.26 | 9.14 | 30.41 | MR |
| **GB2p** | 94.94 | 76.04 | 83.89 | 83.12 | HR |
| **GB3c** | 0.00 | 0.00 | 0.00 | 0.00 | **NR** |
| **GB3p** | 19.76 | 0.00 | 9.14 | 7.87 | LR |
| **GB4c** | 0.00 | 0.00 | 0.01 | 0.01 | **NR** |
| **GB4p** | 27.32 | 38.76 | 0.00 | 17.47 | LR |
| **GB5** | 0.00 | 0.00 | 0.70 | 0.35 | **NR** |
| **GB6c** | 0.34 | 0.00 | 0.00 | 0.06 | **NR** |
| **GB6p** | 0.00 | 0.00 | 0.00 | 0.00 | **NR** |
| **GB7c** | 0.00 | 0.00 | 3.15 | 1.58 | **NR** |
| **GB7p** | 17.89 | 0.00 | 0.34 | 3.15 | **NR** |
| **GB8** | 0.07 | 17.46 | 62.24 | 36.95 | MR |
| **GB9** | 45.15 | 0.16 | 66.72 | 40.94 | MR |
| **GB10** | 0.00 | 0.00 | 0.00 | 0.00 | NR |
| **GB11** | 21.36 | 0.00 | 0.00 | 3.56 | **NR** |
| **GB12** | 0.00 | 0.00 | 0.00 | 0.00 | **NR** |
| **GB13** | 6.32 | 72.94 | 42.14 | 46.44 | MR |
| **GB14** | 0.00 | 0.00 | 13.96 | 6.98 | LR |
| **GB15** | 59.33 | 34.37 | 90.89 | 66.79 | HR |
| **GB16** | 0.95 | 49.95 | 72.73 | 53.17 | HR |

**Table S6** Mean expression mRNA levels (FPKM) of 42 differentially expressed genes are reported in Non-Resp (n=9) and Resp groups (N=9)**.** P-values from CuffDiff analysis were reported. The 17 gene signature panel is highlighted in light blue.

| GENES | DEG (FPKM) | | | | GENES | DEG (FPKM) | | | |
|---|---|---|---|---|---|---|---|---|---|
| | RESP Mean±DS | NON RESP Mean±DS | Log2Fold | p_value | | RESP Mean±DS | NON RESP Mean±DS | Log2Forld | p_value |
| **ABCG2** | 13.46±9.82 | 3.06±3.34 | -2.10 | 0.0015 | **KCNJ10** | 29.36±17.42 | 10.75±16.21 | -1.44 | 0.0021 |
| **ALPK2** | 1.27±1.97 | 4.02±3.61 | 1.59 | 0.0009 | **LGI4** | 11.36±12.76 | 1.62±1.07 | -2.74 | 0.00005 |
| **ANKRD28** | 13.33±3.01 | 0±0 | -7.07 | 1.00 | **MDM1** | 7.69±2.41 | 84.04±4.32 | 3.43 | 0.00005 |
| **BIRC3** | 4.25±3.6 | 12.1±11.06 | 1.49 | 0.00055 | **MEOX2** | 15.79±10.2 | 4.8±4.04 | -1.70 | 0.0038 |
| **CA3** | 4.41±4.19 | 29.61±39.53 | 2.72 | 0.00005 | **NDRG2** | 140.7±140.76 | 33.44±27.09 | -2.07 | 0.00075 |
| **CA9** | 5.28±6.98 | 18.47±21.25 | 1.79 | 0.00235 | **NLGN3** | 35.7±20.45 | 12.04±9.62 | -1.56 | 0.0018 |
| **CAV1** | 30.2±18.04 | 126.81±134.4 | 2.07 | 0.00055 | **NNMT** | 28.28±33.53 | 95.01±75.29 | 1.74 | 0.00225 |
| **CDH4** | 13.9±11.55 | 4.14±1.97 | -1.72 | 0.00075 | **PBX3** | 8.84±3.29 | 23.9±16.27 | 1.42 | 0.0016 |
| **CLDN5** | 38.65±29.45 | 12.37±12.37 | -1.64 | 0.0037 | **PCSK6** | 18.22±44.44 | 3.55±3.98 | -2.33 | 0.0013 |
| **COL8A2** | 5.96±4.75 | 18.42±25.64 | 1.61 | 0.001 | **PLP1** | [illegible]47.25±1345.[illegible] | [illegible]70.27±137.53 | -3.20 | 0.00005 |
| **CXCL14** | 13.44±10.02 | 41.41±34.9 | 1.62 | 0.00015 | **PRICKLE1** | 3±0.92 | 12.07±12.54 | 1.97 | 0.00125 |
| **EDNRB** | 51.99±44.54 | 16.83±12.37 | -1.62 | 0.00295 | **PTPRD** | 18.25±12.83 | 0±0 | -7.52 | 1 |
| **EGFR** | [illegible]21.93±614.5[illegible] | [illegible]51.82±312.1[illegible] | -2.03 | 0.0042 | **RAB27B** | 1.19±0.93 | 4.85±5.33 | 1.94 | 0.00145 |
| **EGFR-AS1** | 61.64±69.68 | 6.9±10.52 | -3.14 | 0.0011 | **RHOJ** | 31.23±24.23 | 12.57±5.81 | -1.31 | 0.0005 |
| **ENPP5** | 3.37±4.11 | 11.46±18.62 | 1.74 | 0.00105 | **SDC1** | 3.99±1.7 | 11.8±9.37 | 1.54 | 0.0017 |
| **F5** | 0.58±0.65 | 2.85±3.87 | 2.12 | 0.00015 | **SDPR** | 9.53±8.77 | 2.32±0.84 | -1.99 | 0.00045 |
| **FGFR3** | 39.04±27.95 | 9.17±11.13 | -2.08 | 0.00015 | **SEMA3A** | 2.62±2.94 | 11.41±15.42 | 2.08 | 0.0004 |
| **FOXG1** | 44.88±38.63 | 11.25±10.7 | -1.99 | 0.00175 | **SEMA6A** | 30.77±16.69 | 13.47±6.44 | -1.19 | 0.00305 |
| **GEM** | 12.13±6.18 | 33.93±23.46 | 1.48 | 0.0009 | **SH3GL2** | 2.16±2.48 | 7.37±11.42 | 1.72 | 0.0008 |
| **GLIPR1** | 13.07±5.94 | 45.64±36.22 | 1.80 | 0.003 | **TRPM3** | 11.06±13.64 | 2.63±1.36 | -2.03 | 0.00285 |
| **GRM3** | 5.95±7.09 | 1.77±1.75 | -1.69 | 0.0036 | **UNC5D** | 3.19±5.23 | 0.85±0.75 | -1.79 | 0.0002 |

**Table S7** GB cases molecular subtypes classification based on TCGA gene expression profile of 490 genes (Anderson Cancer Center)

| GB cases | Predicted GB Molecular Subtype | Prob(Pred) | Others | Phasor FLIM outcome | %DR |
|---|---|---|---|---|---|
| GB2c | Proneural | 0.945718702 | | Resp | MR |
| GB3c | Mesenchymal | 0.988689492 | | Non-Resp | NR |
| GB3p | Mesenchymal | 0.991351319 | | Resp | LR |
| GB4c | Mesenchymal | 0.856475148 | Classical 0.11 | Non-Resp | NR |
| GB4p | Mesenchymal | 0.826027351 | | Resp | LR |
| GB6c | Mesenchymal | 0.775490974 | Neural 0.13 | Non-Resp | NR |
| GB6p | Mesenchymal | 0.864154868 | Proneural 0.11 | Non-Resp | NR |
| GB7c | G-CIMP | 0.997111142 | | Non-Resp | NR |
| GB7p | Mesenchymal | 0.832116053 | Classical 0.12 | Non-Resp | NR |
| GB8 | Classical | 0.988673996 | | Resp | MR |
| GB9 | Neural | 0.532155727 | Classical 0.34 Mesenchymal 0.12 | Resp | MR |
| GB10 | Classical | 0.597423274 | Mesenchymal 0.21 Neural 0.17 | Non-resp | NR |
| GB11 | Mesenchymal | 0.996821433 | | Non-resp | NR |
| GB12 | Mesenchymal | 0.995641967 | | Non-resp | NR |
| GB13 | Classical | 0.780092979 | Neural 0.14 | Resp | MR |
| Gb14 | Classical | 0.610788371 | Mesenchymal 0.31 | Resp | LR |
| GB15 | Classical | 0.646330242 | Neural 0.28 | Resp | HR |
| GB16 | Neural | 0.569172723 | Classical 0.37 | Resp | HR |
| **Core and Periphery** | | | | | |
| GB3c | Mesenchymal | 0.988689492 | | Non-Resp | NR |
| GB3p | Mesenchymal | 0.991351319 | | Resp | LR |
| GB4c | Mesenchymal | 0.856475148 | Classical 0.11 | Non-Resp | NR |
| GB4p | Mesenchymal | 0.826027351 | | Resp | LR |
| GB6c | Mesenchymal | 0.775490974 | Neural 0.13 | Non-Resp | NR |
| GB6p | Mesenchymal | 0.864154868 | Proneural 0.11 | Non-Resp | NR |
| GB7c | G-CIMP | 0.997111142 | | Non-Resp | NR |
| GB7p | Mesenchymal | 0.832116053 | Classical 0.12 | Non-Resp | c |

**Table S8** Somatic mutations in genes known to be altered in IDH1 WT GB. Group, Resp, Responder tumors, Non-resp, Non-responder tumors. Sample, tumor in which the mutation has been identified. Gene, mutated gene. Chr, Chromosome location of the gene. Start, chromosomal position where the variant begins. End, chromosomal position where the variant ends. Ref Allele, wild type reference allele. Tum Allele, tumor allele changed by mutation. Variant classification, type of mutation. Protein change, amino acid change

| Group | Sample | Gene | Chr | Start | End | Ref Allele | Tum Allele | Variant Classificatio | Protein Change |
|---|---|---|---|---|---|---|---|---|---|
| Resp | GB9 | TP53 | 17 | 7577091 | 7577091 | G | A | Missense | p.Arg283Cys |
| Non-resp | GB3C | RET | 10 | 43609920 | 43609920 | A | C | Splice Site | |
| Non-resp | GB4c | RET | 10 | 43609920 | 43609920 | A | C | Splice Site | |
| Resp | GB15 | RET | 10 | 43609920 | 43609920 | A | C | Splice Site | |
| Non-resp | GB6c | RB1 | 13 | 48942685 | 48942685 | C | T | Nonsense | p.Arg358Ter |
| Resp | GB2c | PTPN11 | 12 | 1,13E+08 | 1,13E+08 | G | T | Missense | p.Gly503Val |
| Non-resp | GB12 | PTPN11 | 12 | 1,13E+08 | 1,13E+08 | G | A | Missense | p.Ala461Thr |
| Resp | GB2c | PTEN | 10 | 89692769 | 89692769 | G | A | Splice Site | |
| Non-resp | GB3c | PTEN | 10 | 89720721 | 89720721 | - | A | Frame Shift In | p.Asn292LysfsTer6 |
| Non-resp | GB5 | PTEN | 10 | 89685269 | 89685269 | G | A | Splice Site | |
| Non-resp | GB1 | PTEN | 10 | 89720726 | 89720726 | G | - | Frame Shift De | p.Gly293GlufsTer14 |
| Non-resp | GB6c | PIK3R1 | 5 | 67589149 | 67589149 | A | C | Missense | p.Lys379Asn |
| Resp | GB16 | PIK3CA | 3 | 1,79E+08 | 1,79E+08 | A | T | Missense | p.Glu542Val |
| Non-resp | GB12 | NOTCH1 | 9 | 1,39E+08 | 1,39E+08 | C | G | Splice Site | |
| Resp | GB15 | NOTCH1 | 9 | 1,39E+08 | 1,39E+08 | C | G | Splice Site | |
| Non-resp | GB5 | NF1 | 17 | 29527438 | 29527438 | - | A | Splice Site | |
| Non-resp | GB5 | NF1 | 17 | 29557285 | 29557285 | C | - | Frame Shift De | p.Arg1000ValfsTer12 |
| Non-resp | GB7c | NF1 | 17 | 29497001 | 29497001 | A | C | Missense | p.Lys191Thr |
| Non-resp | GB12 | NF1 | 17 | 29527461 | 29527461 | C | T | Nonsense | p.Arg304Ter |
| Resp | GB14 | NF1 | 17 | 29497001 | 29497001 | A | C | Missense | p.Lys191Thr |
| Non-resp | GB1 | NF1 | 17 | 29585420 | 29585420 | T | G | Missense | p.Leu1411Arg |
| Non-resp | GB10 | MED12 | 23 | 70349253 | 70349253 | C | G | Missense | p.Ala1222Gly |
| Non-resp | GB12 | MED12 | 23 | 70346203 | 70346203 | G | T | Missense | p.Val852Phe |
| Resp | GB2C | EGFR | 7 | 55259422 | 55259422 | A | T | Missense | p.Tyr827Phe |
| Resp | GB14 | EGFR | 7 | 55233043 | 55233043 | G | T | Missense | p.Gly598Val |
| Resp | GB16 | EGFR | 7 | 55221716 | 55221716 | T | A | Missense | p.Phe254Ile |
| Resp | GB16 | CDKN2A | 9 | 21971142 | 21971142 | G | T | Nonsense | p.Cys72Ter |
| Non-resp | GB3c | CREBBP | 16 | 3786721 | 3786721 | T | G | Missense | p.Lys1497Thr |
| Non-resp | GB4c | CREBBP | 16 | 3786740 | 3786740 | G | T | Missense | p.Gln1491Lys |
| Non-resp | GB5 | CREBBP | 16 | 3786733 | 3786733 | A | T | Missense | p.Ile1493Lys |
| Resp | GB9 | CREBBP | 16 | 3786721 | 3786721 | T | G | Missense | p.Lys1497Thr |
| Resp | GB9 | CREBBP | 16 | 3842060 | 3842060 | G | T | Missense | p.His418Asn |
| Non-resp | GB10 | CREBBP | 16 | 3786721 | 3786721 | T | G | Missense | p.Lys1497Thr |
| Non-resp | GB10 | CREBBP | 16 | 3786733 | 3786733 | A | T | Missense | p.Ile1493Lys |
| Non-resp | GB12 | CREBBP | 16 | 3786721 | 3786721 | T | G | Missense | p.Lys1497Thr |
| Resp | GB13 | CREBBP | 16 | 3786721 | 3786721 | T | G | Missense | p.Lys1497Thr |
| Resp | GB14 | CREBBP | 16 | 3786721 | 3786721 | T | G | Missense | p.Lys1497Thr |
| Resp | GB15 | CREBBP | 16 | 3786721 | 3786721 | T | G | Missense | p.Lys1497Thr |
| Resp | GB16 | CREBBP | 16 | 3786721 | 3786721 | T | G | Missense | p.Lys1497Thr |
| Non-resp | GB12 | ARID1A | 1 | 27100950 | 27100950 | A | C | Missense | p.Gln1411Pro |
| Non-resp | GB12 | ARID1A | 1 | 27100958 | 27100958 | A | C | Missense | p.Ser1414Arg |
| Resp | GB14 | ARID1A | 1 | 27100950 | 27100950 | A | C | Missense | p.Gln1411Pro |
| Resp | GB14 | A2M | 12 | 9256992 | 9256992 | C | T | Missense | p.Arg370His |

**Table S9** Statistically significant genes that differentiate the two groups Non-responder and Responder. Group, Resp, Responder tumors, Non-resp, Non-responder tumors. Sample, tumor in which the mutation has been identified. Gene, mutated gene. Chr, Chromosome location of the gene. Position, chromosomal position where the variant occurs. Type, type of mutation. Ref Allele, wild type reference allele. Tum Allele, tumor allele changed by mutation. Variant, type of mutation. Protein change, amino acid change. Each variant is shown in detail. Two variants of FBN3, $FBN3^{E492K}$ in GB14 and $FBN3^{R2688Q}$ in GB16, were already described and annotated in COSMIC with the IDs COSM9337963 and COSM3541504, respectively. The impact on protein function and thus clinical significance has not yet been annotated for most of these variants resulting in "unknown significance" for the Varsome classification. Only the two splicing variants in DNAH5 in sample GB6c and in DENND6B in sample GB13 are predicted to be pathogenic. Three genes (ZNF713, DNAH5 and SLC7A4) were mutated only in the Non-Resp group with a percentage of 67% (6/9), 56% (5/9) and 56% (5/9), respectively. GALNTL5, FBN3 and DENND6B were shared only by the Resp group with a frequency of 56%. The OTOR gene was in common between groups of tumors, but with higher mutated rate in Resp cases. To the best of our knowledge none of the genes found have ever been associated to glioblastoma.

| Group | Sample | Gene | Chr | Position | Ref All | Tum All | Variant | Protein Change |
|---|---|---|---|---|---|---|---|---|
| Non-resp | GB4c | ZNF713 | 7 | 55991292 | GT | AC | Splice Site | |
| Non-resp | GB5 | ZNF713 | 7 | 55991292 | GT | AC | Splice Site | |
| Non-resp | GB6c | ZNF713 | 7 | 55991292 | GT | AC | Splice Site | |
| Non-resp | GB7c | ZNF713 | 7 | 55991292 | GT | AC | Splice Site | |
| Non-resp | GB12 | ZNF713 | 7 | 55991292 | GT | AC | Splice Site | |
| Non-resp | GB1 | ZNF713 | 7 | 55991292 | GT | AC | Splice Site | |
| Resp | GB2c | GALNTL5 | 7 | 151716807 | A | C | ssense Mutat | p.Lys418Thr |
| Resp | GB9 | GALNTL5 | 7 | 151716807 | A | C | ssense Mutat | p.Lys418Thr |
| Resp | GB13 | GALNTL5 | 7 | 151716807 | A | C | ssense Mutat | p.Lys418Thr |
| Resp | GB14 | GALNTL5 | 7 | 151716807 | A | C | ssense Mutat | p.Lys418Thr |
| Resp | GB9 | FBN3 | 19 | 8150340 | A | C | ssense Mutat | p.Trp2332Gly |
| Resp | GB13 | FBN3 | 19 | 8150340 | A | C | ssense Mutat | p.Trp2332Gly |
| Resp | GB14 | FBN3 | 19 | 8200962 | C | T | ssense Mutat | p.Glu492Lys |
| Resp | GB16 | FBN3 | 19 | 8136957 | C | T | ssense Mutat | p.Arg2688Gln |
| Resp | GB2c | DENND6B | 22 | 50752708 | G | C | Splice Site | |
| Resp | GB9 | DENND6B | 22 | 50752708 | G | C | Splice Site | |
| Resp | GB13 | DENND6B | 22 | 50752703 | T | A | Splice Site | |
| Resp | GB14 | DENND6B | 22 | 50752708 | G | C | Splice Site | |
| Non-resp | GB3c | SLC7A4 | 22 | 21384444 | G | C | ssense Mutat | p.Phe393Leu |
| Non-resp | GB4c | SLC7A4 | 22 | 21384444 | G | C | ssense Mutat | p.Phe393Leu |
| Non-resp | GB5 | SLC7A4 | 22 | 21384444 | G | C | ssense Mutat | p.Phe393Leu |
| Non-resp | GB6c | SLC7A4 | 22 | 21384444 | G | C | ssense Mutat | p.Phe393Leu |
| Non-resp | GB10 | SLC7A4 | 22 | 21384444 | G | C | ssense Mutat | p.Phe393Leu |
| Non-resp | GB5 | DNAH5 | 5 | 13868104 | AA | - | Splice Site | |
| Non-resp | GB6c | DNAH5 | 5 | 13845088 | T | G | ssense Mutat | p.Lys1710Thr |
| Non-resp | GB6c | DNAH5 | 5 | 13845095 | C | G | ssense Mutat | p.Glu1708Gln |
| Non-resp | GB6c | DNAH5 | 5 | 13845100 | T | A | ssense Mutat | p.Tyr1706Phe |
| Non-resp | GB6c | DNAH5 | 5 | 13845102 | C | A | Splice Site | c.5115G>Tp.Gly1705= |
| Non-resp | GB6c | DNAH5 | 5 | 13845104 | T | A | Splice Site | |
| Non-resp | GB12 | DNAH5 | 5 | 13845088 | T | G | ssense Mutat | p.Lys1710Thr |
| Non-resp | GB12 | DNAH5 | 5 | 13717589 | C | A | ssense Mutat | p.Gln4180His |
| Non-resp | GB1 | DNAH5 | 5 | 13794173 | - | A | Splice Site | |
| Non-resp | GB4c | OTOR | 20 | 16730588 | A | T | ssense Mutat | p.Tyr99Phe |
| Resp | GB8 | OTOR | 20 | 16730588 | A | T | ssense Mutat | p.Tyr99Phe |
| Resp | GB9 | OTOR | 20 | 16730588 | A | T | ssense Mutat | p.Tyr99Phe |
| Resp | GB13 | OTOR | 20 | 16730588 | A | T | ssense Mutat | p.Tyr99Phe |
| Resp | GB14 | OTOR | 20 | 16730588 | A | T | ssense Mutat | p.Tyr99Phe |
| Resp | GB15 | OTOR | 20 | 16730588 | A | T | ssense Mutat | p.Tyr99Phe |

**Video S1** Video shooting (10X magnification, 9 tiles assembly) of a GB-EXP over-night. At 2-week time, the vitality of GB-EXPs cultured in matrigel was determined by setting up over-night live-imaging analyses which revealed an intensive cell activity particularly evident at the surface of the explant in contact with the surrounding cells and neighboring explant.

GB-EXPs VIDEO S1.avi
